# Adsorption of Azo Dyes by a Novel Bio-Nanocomposite Based on Whey Protein Nanofibrils and Nano-clay: Equilibrium Isotherm and Kinetic Modeling

**DOI:** 10.1101/2020.11.24.394205

**Authors:** ShabBoo Rahimi Aqdam, Daniel E. Otzen, Niyaz Mohammad Mahmoodi, Dina Morshedi

**Affiliations:** Bioprocess Engineering Research Group, Industrial and Environmental Biotechnology, National Institute of Genetic Engineering and Biotechnology, Tehran, Iran; Interdisciplinary Nanoscience Centre (iNANO), Department of Molecular Biology and Genetics, Aarhus University, 8000 Aarhus C, Denmark; Department of Environmental Research, Institute for Color Science and Technology, Tehran, Iran

**Author notes:** Corresponding author: D. Morshedi.

**Keywords:** Protein-based nanocomposite, Whey Protein Concentrate (WPC), Montmorillonite, Chrysoidine, Wastewater remediation, Dye pollution

## Abstract

Excessive discharge of hazardous azo dyes into the aquatic ecosystem is a global environmental concern. Here, we develop a green approach to remediate dye pollutions in water by fabricating an easy-separable bio-nanocomposite, based on whey protein concentrate, its nanofibrils, and montmorillonite nano-clay. To characterize the nanocomposite, we used SEM, FT-IR, XRD, and BET techniques. Nanofibrils lead to a uniform dispersion of montmorillonite in the whey protein matrix and also reinforce the nanocomposite. The adsorption efficacy was monitored in a batch system, using cationic dyes (Chrysoidine-G, Bismarck brown-R), reactive dyes (reactive black-5, reactive orange-16), acid dyes (acid red-88, acid red-114), and direct dyes (direct violet-51, Congo red). This nanocomposite adsorbed different dye classes, cationic dyes quicker (> 82%, after 4 h), and reactive dyes slower. Then, the effect of initial dye concentration, pH, contact time, adsorbent dose, and temperature on Chrysoidine-G adsorption was explored. The adsorbent showed a high removal (>93%) for a wide concentration range of Chrysoidine-G, also acidic pH and higher temperature are more favorable for the process. Equilibrium adsorption parameters were reasonably fitted with a linear (Nernst) isotherm model. The results indicated the existence of an unlimited number of absorption sites, *i.e.* no saturation was achieved under our experimental conditions (q_max(Exp)_= 731 mg/g). Kinetic data were fitted with pseudo-second-order and intra-particle diffusion models. We conclude that this nanocomposite is a green adsorbent with potential use for wastewater treatment and related purposes.

**Highlights:**

- We produced an easy-separable bio-nanocomposite using whey nanofibrils and MMT, with high adsorption capacity
- Nanofibrils help disperse MMT particles uniformly in the WP matrix
- The adsorbent’s performance was compared to the adsorbents in absence of MMT and nanofibrils
- This composite adsorbs cationic, anionic, direct and reactive azo dyes with different kinetics
- Adsorption isotherms and kinetics are studied in detail

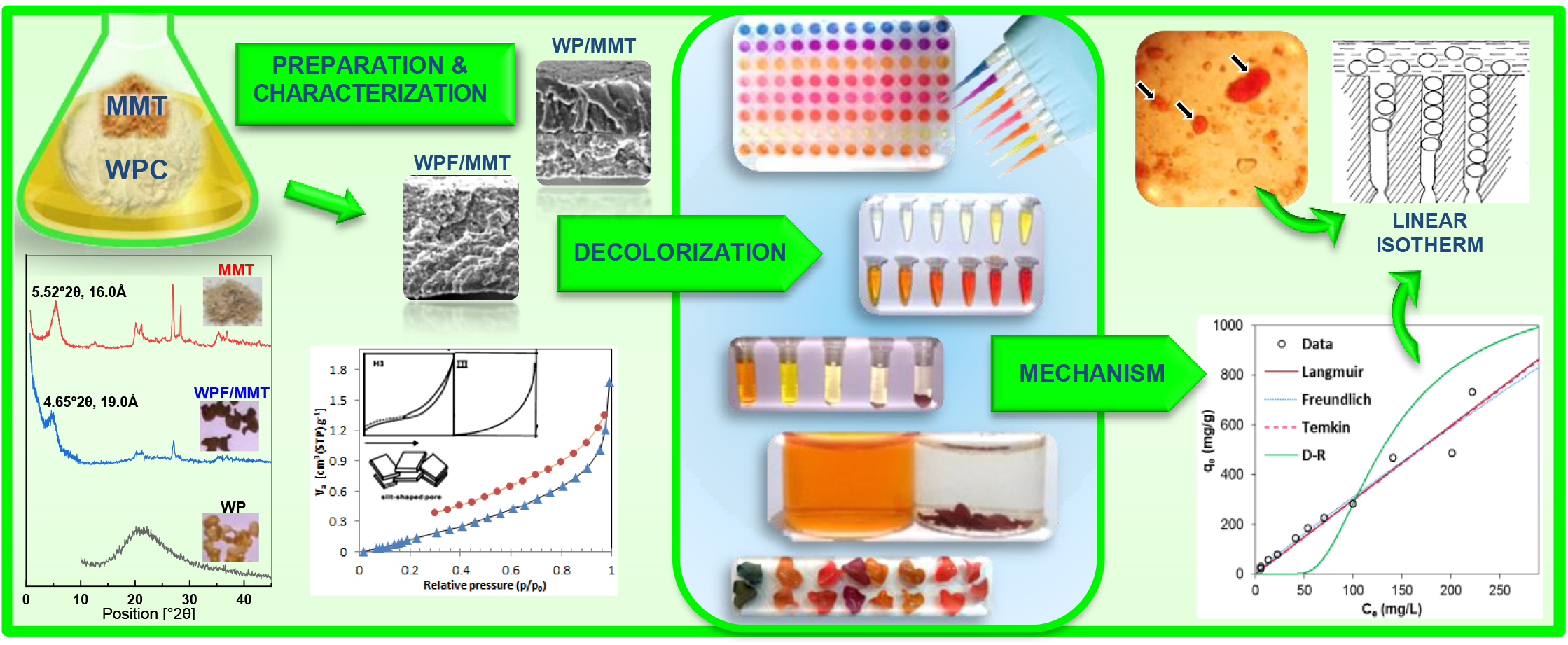

## 1. Introduction

The release of large amounts of unprocessed wastewater into the environment is a global issue with significant health and environmental impacts. For instance, 10-15% of dyes used in dyeing processes are discharged into effluent streams [1]. Azo dyes, because of their cheap and straightforward synthesis, account for 60-70% of total dye production in the world [1]. However, due to their complex structures and aromatic content, they are not naturally degradable [2]; consequently, they constitute a major source of aqueous pollutants worldwide. Besides changing the appearance and color of water, they inhibit photosynthesis through the absorption of sunlight and also lead to carcinogenic and mutagenic metabolites [3, 4].

Wastewater treatment generally includes biological, chemical, and physical methods. Common remediation techniques for dyes include electrochemical destruction, filtration, sedimentation, coagulation and flocculation, ion exchange, adsorption, exposure to light, and microbiological treatment [5–9]. Adsorption techniques are most widespread because of their low cost, high efficiency, simplicity, and usual absence of by-products. However, nano- and micro-scale adsorbents need extra procedures, *e.g.* centrifugation and filtration, to be separated from the solution after the adsorption process; otherwise, they would cause secondary pollution [10, 11]. Meanwhile, due to the vast diversity of the chemo-physical and structural properties of azo dyes, there are no stand-alone methods to decolorize wastewater completely [12, 13]. Moreover, the growth of industries with dye by-product pollution coupled with shrinking water resources makes it imperative to develop new methods and materials to recycle wastewater.

In this regard, natural materials, bio-polymers, and bio-nanocomposites have received much attention as potential adsorbents. Natural materials are environmentally friendly as they are biodegradable and biocompatible. Thanks to their structural and chemical diversity, protein-based adsorbents have distinct advantages over conventional polysaccharide-based adsorbents such as cellulose, chitosan, and lignin [14, 15]. Additionally, some proteins can self-assemble and convert into highly ordered nanofibrils which resist elevated temperature and salt concentrations and also show strength and stiffness comparable to that of steel and silk [16, 17].

Whey protein is a well-known source of protein that easily self-assembles into nanofibrils. Whey is a byproduct of cheese manufacturing, with roughly 50 million tons of unused whey per year [18]. Thanks to whey’s high carbohydrate and protein content, its release as waste into the environment has a significant negative impact due to eutrophication [19–21]. Thus, exploitation of whey can both mobilize unwanted waste material and generate novel functionalities. It is particularly useful that whey can form gels, hydrogels, and also proteinaceous nanofibrils [22]. Such nanofibrils can reinforce composite mixtures by increasing gel viscosity and strength [23]. On top of this, proteinaceous nanofibrils show excellent ability to adsorb azo dyes [24], arsenic [25], organic contaminants [26], and lead (II) [27] from aqueous solutions. Another dual-purpose (*i.e.* promoting both structural strength and adsorption) material is montmorillonite nano-clay (MMT) [28], whose basic structure is a central alumina octahedral sheet sandwiched between two tetrahedral silica sheets. These ~1 nm aluminosilicate layers can be extended indefinitely in the x-y plane. The layers are stabilized by electrostatic and Van der Waals forces with a fixed interlayer distance, which varies depending on the cation. The cations can be exchanged with organic cations (such as dye molecules), leading to increased interlayer distance (d-spacing). Since the layers are overall negatively charged, they reversibly bind cationic dyes [29–31].

Nevertheless, both MMT and nanofibrils are in nano-scale, consequently, their recovery after adsorption of pollutant need some eco-unfriendly extra steps. To address this issue, and also to exploit and combine the advantages of nanofibrils and MMT, we have produced an easily separable bio-nanocomposite (WPF/MMT) based on nanofibrils of whey protein concentrate (WPC) together with MMT. Different classes of azo dyes’ adsorption rate and the effect of contact time on adsorption were investigated using three adsorbents, *i.e.* whey protein polymer (WP), WP composite with MMT but without nanofibrils (WP/MMT), and WP composite combining amyloid nanofibrils and MMT (WPF/MMT). The studied dyes include two cationic dyes (Chrysoidine-G and Bismarck brown-R), two reactive dyes (reactive black 5 and reactive orange 16), two acid dyes (acid red 88 and acid red 114), and also two direct dyes (direct violet 51 and Congo red) (see structures in **Supp. Table 1**). To gain a better insight into the adsorption process, the effect of initial dye concentration, pH, temperature, and adsorbent dose on Chrysoidine-G removal were examined using WPF/MMT nanocomposite. Finally, the adsorption isotherms and kinetics of Chrysoidine-G were studied in detail.

## 2. Materials and Methods

### 2.1. Materials

Whey Protein Concentrate 8010 (WPC with 80% protein) was obtained from Hilmar™ (North Lander Avenue, P.O. Box 910 Hilmar, CA). The montmorillonite K-10 (surface area 220-270 m^2^/g), all 8 azo dyes (**Supp. Table 1**), and Thioflavin-T (ThT) were from Sigma-Aldrich (St. Louis, MO). Glycerol, all salts, and other materials were from Merck (Darmstadt, Germany).

### 2.2. Preparation of WPC Polymer and Nanocomposites

To prepare MMT suspensions, 6 g of MMT was added to 20 mL deionized water in a sealed beaker and was heated (~80#x00B0;C) with continuous magnetic stirring for 1 h, followed by sonication for 30 min in an ultrasonic bath (JAC 2010 KODO, Gyeonggi-do, Korea) to disperse MMT. **Fig. 1** illustrates the key steps for the preparation of different whey protein-based polymer and composites, without nanofibrils (the black-arrows path), and with nanofibrils (the white-arrows path).

**Figure 1:**
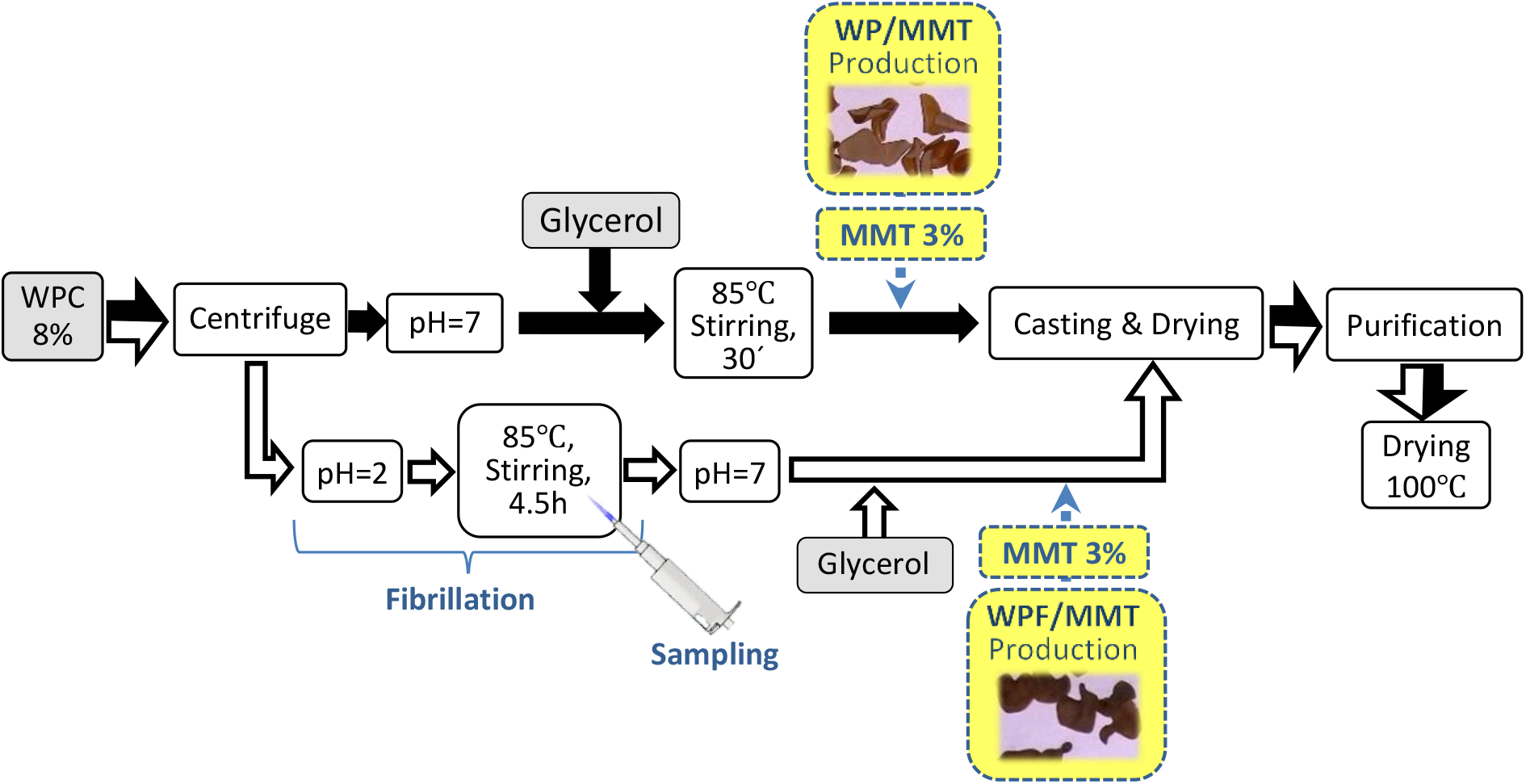
Main steps of WP polymer and nanocomposites production. The black-arrows show WP and WP/MMT production steps; while to produce WPF and WPF/MMT the white-arrows should be followed. Dispersed MMT should be added before the casting step to produce WP/MMT and WPF/MMT nanocomposites,. The black and white arrows show the common steps between the two paths.

In both pathways, nanocomposites are formed by adding 3% w/v of dispersed MMT to the film-forming solution before casting. At the first stage, 8% (w/v) of WPC was hydrated in deionized water using a magnetic stirrer, and centrifuged (13000 rpm, 30 min, at 4°C) to remove insoluble protein [32]. To produce films without nanofibrils (WP and WP/MMT), the pH was adjusted to 7 using NaOH (6M), glycerol (40% v/w WPC) was added, and the solution was stirred at 85°C for 30 min in a water bath. To produce films containing nanofibrils (WPF and WPF/MMT), the pH was adjusted to 2 using HCl (12M), and the solution stirred at 85°C for 4.5 h [33, 34]. During this step, sampling was performed to monitor the fibrillation process (see section 3.1). The resulting solution was then cooled, and the pH was adjusted to 7. Subsequently glycerol (40% v/w WPC), and for the WPF/MMT nanocomposite, 3% dispersed MMT, were added, and the mixtures were stirred thoroughly before casting. Afterward, the cast solution was dried at 37°C overnight, followed by 2 h at 70°C. Finally, the films were immersed in distilled water at 4°C for 48 h to remove non-cross-linked parts [35] and then dried at 100°C for 1 h.

## 3. Assessment of Nanofibril Formation

### 3.1. ThT Fluorescence Assay

We used ThT fluorescence to monitor the progress of fibrillation [36, 37]. At different times during fibrillation (**Fig. 1**), a 10 μL sample of the solution was added to 490 μL of 12 μM ThT solution in Tris buffer 10mM (pH 8). Subsequently, the fluorescence emission spectra of samples were recorded in the range of 450 to 550 nm with excitation at 440 nm (Varian Cary Eclipse fluorescence spectrophotometer, Australia). The excitation and emission slit widths were 5 and 10 nm, respectively.

### 3.2. Transmission Electron Microscopy (TEM)

After fibrillation and adjusting the pH to 7 (**Fig. 1**), 5 μL of the solution was placed onto a carbon-coated, glow-discharged 400-mesh grid for 30 s. The grid was stained with phosphotungstic acid 1% (pH 6.8) and blotted dry. Images were recorded on a TEM microscope (JEM-1010; JEOL, Tokyo, Japan) at 60 kV, using an Olympus KeenView G2 camera.

### 3.3. Material Characterization

#### 3.3.1. Scanning Electron Microscopy (SEM)

The morphology of the surfaces and fractured cross-sections of all films were visualized using SEM (VEGAII TESCAN, Czech Republic). To prepare the fractured cross-section, pieces of the WP films were frozen by immersing in liquid nitrogen and then fractured manually. The surface and fracture sides of the films were mounted on the specimen stubs and then sputter-coated with a thin layer of gold and placed into the scanning electron microscope to observe the surface and cross-section morphology of the films.

#### 3.3.2. Fourier-Transform Infrared (FT-IR) Spectroscopy

FT-IR spectra of the WPF/MMT, WP, and pristine MMT were recorded using a Perkin-Elmer Spectrum One spectrometer (Waltham, MA, USA) at the range of 400-4000 cm^−1^. Powdered samples were mixed and ground with KBr powder to make packed tablets for FT-IR measurement.

#### 3.3.3. X-ray Diffraction (XRD)

XRD measurements were performed on the WPF/MMT, WP, and pristine MMT using a Philips PW 1730 diffractometer (Eindhoven, The Netherlands), employing Cu Kα radiation source (λ= 1.54060 Å). The data were collected for 2θ values 0.71-9.99° in 0.02° step size, and 10.00-79.95 values in 0.05° step sizes.

#### 3.3.4. Nitrogen Adsorption Isotherm (BET)

Nitrogen adsorption/desorption isotherms were measured with a BELSORP-miniII (Osaka, Japan) Brunauer-Emmett-Teller (BET) analyzer at 77 K. Before measurements, the nanocomposite samples were degassed at 180°C in a vacuum chamber for 8 h.

### 3.4. Batch Dye Adsorption Studies

We first investigated the decolorization percentage of WPF/MMT compared with WP, WP/MMT by immersing 2% w/v adsorbent in the dye solutions. The initial concentration of the dyes in distilled water was 250 mg/L. During the decolorization process, 200 μL of the dye solutions were collected at different times and their absorbance was measured using an EPOCH12 plate reader (BioTek, Winooski, Vermont, USA) at their maximum absorbance wavelength (**Supp. Table 1**). The percentage of decolorization (%) was calculated as follows:

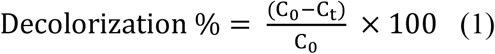

where C_0_ and C_t_ (mg/L) are the concentrations of the examined dyes at the beginning of the experiment and after time t, respectively.

The effect of initial dye pH (2-10) and temperature (4-45°C) on adsorption of Chrysoidine (100 mg/L) was studied using 9 g/L of WPF/MMT. Furthermore, the influence of different concentrations of Chrysoidine-G (25-250 mg/L) and adsorbent dosage (1.45-17.3 g/L) were carried out in a reaction volume of 1.5 mL in 96-deep-well plates (Extragene, Taichung, Taiwan), which were covered with adhesive plate seals (Thermo scientific, USA), and put into an orbital shaker with 300 rpm speed (HOLDEKER5000D Orbital motion-UK) at 35°C. At different time intervals q_t_, the amount of adsorbed dye per gram of adsorbent (mg/g), was measured as follows:

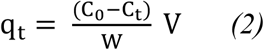

where C_0_ and C_t_ are the concentrations of the examined dyes (in mg/L) at the beginning and after time t, respectively, W is the weight of the adsorbent (g), and V is the volume of solution (L).

Equilibrium parameters for Chrysoidine-G adsorption were analyzed using different WPF/MMT dosages (0.04-12.7 g/L) at 37°C, using linear (Nernst), Longmuir, Freundlich, Temkin, and Dubinin-Radushkevich (D-R) Isotherm models:

#### Linear (Nernst) Isotherm

The linear isotherm is the simplest adsorption isotherm, represented by the following equation:

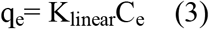

where q_e_ (mg/g) is the amount of adsorbed dye at equilibrium, K_linear_ is the adsorption coefficient (L/g) [38] and C_e_ (mg/L) is the equilibrium concentration of free (not adsorbed) dye in the solution.

#### Langmuir Isotherm

The Langmuir isotherm assumes monolayer formation and homogenous adsorption sites, which can be represented by the following equation:

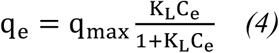

where q_e_ (mg/g) is the amount of adsorbed dye at equilibrium, C_e_ (mg/L) is the equilibrium concentration dye, q_max_ (mg/g) is the maximum amount of adsorbed dye and K_L_ (L/mg) is the Langmuir constant related to the affinity of the surficial binding sites [39].

#### Freundlich Isotherm

The Freundlich Isotherm equation is expressed as follows:

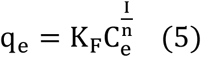

where K_F_ (mg/g) and n are constants for the adsorbate (dye) and adsorbent (nanocomposite),respectively [40].

#### Temkin Isotherm

The Temkin isotherm is expressed as follows [41]:

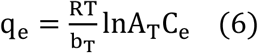

where R is the universal gas constant (8.314 J mol^−1^ K^−1^), *T* is the absolute temperature, b is a constant related to the heat of adsorption (J mol^−1^), and A_T_ is the Temkin isotherm constant (Lg^−1^).

#### D-R Isotherm

The D-R isotherm is given as follows [42]:

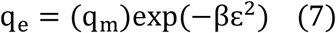

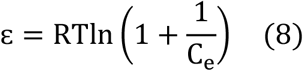

where q_m_ (mg/g) is the D-R monolayer capacity, β is the sorption energy constant and ∊ is the Polanyi potential related to the equilibrium concentration and absolute temperature (T).

Kinetic models of adsorption were assessed using WPF/MMT nanocomposite (8 g/L) in two different concentrations of Chrysoidine-G (50 and 100 mg/L). This experiment was carried out in 10 mL by incubating the samples in the orbital shaker (300 rpm) at 35°C for 48 h. Then, the results were analyzed using pseudo-first-order, pseudo-second-order, and intra-particle diffusion kinetic models.

#### Pseudo-first-order

The linear form of the pseudo-first-order kinetic model [43] is generally expressed as follows:

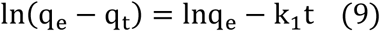

where q_e_ and q_t_ (mg/g) are the amounts of the adsorbed dye at equilibrium and at the measured time respectively and k_1_ is the pseudo-first-order rate constant (1/min) [44].

#### Pseudo-second-order

The linearized form of the pseudo-second-order model [45] is expressed as:

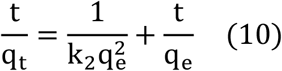

where q_e_ and q_t_ (mg/g) are the amounts of adsorbed dye at equilibrium and at the time t (min), respectively while k_2_ is the pseudo-second-order rate constant of sorption. The plot of t/q_t_ versus t gives a straight line in which q_e_ and k_2_ could be calculated from the slope (1/k_2_q_e_^2^) and intercept (1/q_e_), respectively [44].

#### Intra-particle diffusion

The intra-particle diffusion kinetic model [46] is used to investigate whether intra-particle diffusion is the rate-limiting step in the adsorption of Chrysoidine-G by WPF/MMT. The linear form of the intra-particle diffusion equation can be presented as:

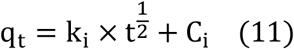

where k_i_ (mg/g min^1/2^) is the intra-particle diffusion rate constant and C_i_ (mg/g) is the intercept of the plot related to the boundary layer influence [47].

## 4. Results and Discussion

### 4.1. Assessment of Nanofibril Formation

Amyloidogenic protein/ peptides, as implied in the name, can self-assemble into highly ordered nanofibrils under particular conditions. According to previous studies [33, 34], we fibrillated whey protein at pH 2 and 85°C and confirmed amyloid formation using the fluorescent amyloid reporter ThT along with TEM images (**Supp. Fig. 1**). ThT fluorescence is a common method to detect amyloid formation *in vitro* [36]. Though not entirely-quantitative, ThT fluorescence intensity is related to the overall mass of the formed fibrils [37]. The increasing fluorescence intensity of ThT over time (**Supp. Fig. 1a**) suggested that WP was undergoing fibril formation. Also, the presence of nanofibrils was confirmed by TEM images (**Supp. Fig. 1b**).

### 4.2. Materials Characterization

#### 4.2.1. SEM Analysis

The microstructure and morphology of the produced materials were studied by SEM (**Fig. 2**).

**Figure 2:**
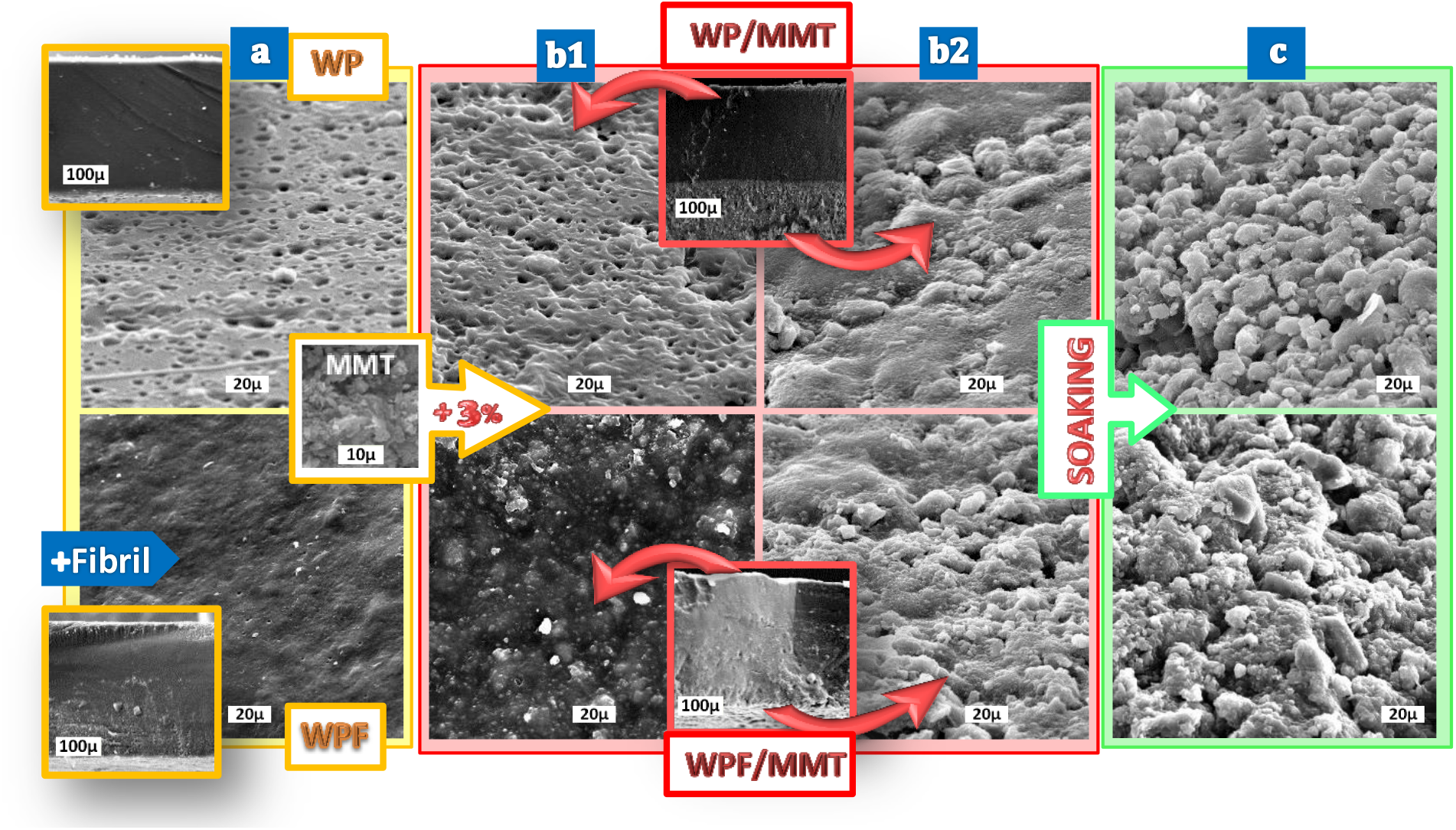
SEM micrographs of the surface (main images) and cross-section (smaller images) of WP polymers and composites with and without nanofibrils (top and bottom row, respectively). Column **(a)** shows WP (top) and WPF (bottom). Columns **(b1, b2)** show upward and downward sides of WP/MMT (top), and WPF/MMT (bottom), **(c)** is their upper surface after soaking in water.

WP showed a rough and porous surface, while its cross-section is smooth and homogenous. In contrast, WPF (with nanofibrils) exhibited a rough and heterogeneous cross-section. These morphologies might be related to the different viscosities of the film-forming solutions. WPF solution was highly viscous, trapping air bubbles and producing a porous cross-section. Nevertheless, the WPF surface was smooth and undulated because of clustered fibrils, similar to observations by Akkermans and coworkers [23]. WP composites with MMT showed different morphologies on their upward and downward sides (in contact with air and the mold respectively) (**Fig. 2b1, b2**). The upward sides were rather similar to the surface of WP films (**Fig. 3a**); while the downward sides show a more rugged surface, in which MMT particles are visible. Interestingly, soaking in water increased the porosity (**Fig. 2c**). WPF/MMT shows a homogenous cross-section. In remarkable contrast, the nanocomposite without nanofibrils (WP/MMT) exhibited a bilayer structure (**Fig. 2b and 3**).

**Figure 3:**
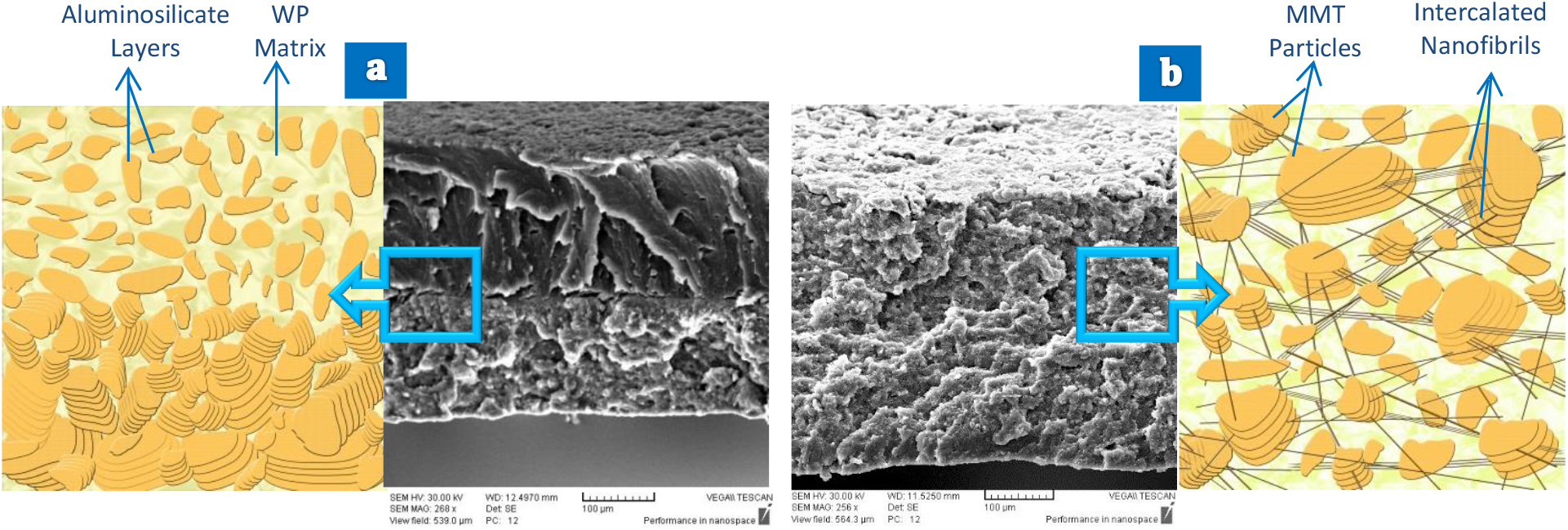
SEM cross-sectional micrographs (the middle images) and the sketches of our hypothesis of arrangement for **(a)** WP/MMT, and **(b)** WPF/MMT nanocomposites. The sketches suggest that in WP/MMT, MMT components settle mostly at the bottom layer. In contrast, they are well dispersed in the WP matrix in the presence of nanofibrils.

By comparing the composites’ cross-sections, we conclude that nanofibrils contribute to the uniform dispersion of MMT particles in the WP matrix. We attribute the bilayer structure to the phase separation between the WP and MMT components due to weak intermolecular forces between these two components. In contrast, the more viscous WPF solution can probably disperse MMT particles better in the matrix due to the crosslinking them by nanofibrils (see sketch in **Fig. 3**). While this bilayer structure has not been reported previously, the structures of WP, WPF, and WPF/MMT are similar to previous reports [21, 48]. The only exception is the work by Kumar et al. (2010) who observe that fracture images are undulated and similar to what we obtained using manually torn samples (**Supp. Fig. 2**) as opposed to fracturing in liquid nitrogen. They noted that “The white strands in the SEM images correspond to MMT platelets” [49, 50]. However, since these strands are present in our WP films too (*i.e.* in the absence of MMT) and we do not observe them in the N_2_-fractured surface, we assume this structure to be caused by the glycerol plasticizer.

#### 4.2.2. XRD Analysis

XRD is a common analysis method to verify composites’ intercalated or exfoliated structures. The XRD pattern of WPF/MMT, WP, and pristine MMT are shown in **Fig. 4a.** The XRD pattern of WP had only a broad peak around 20-22°, indicating a lack of crystallinity and amorphous shape [18]. When MMT was used as filler in the amorphous whey protein matrix, the composite exhibited a similar pattern to the pristine MMT, demonstrating a crystalline form of the composite, which is only modestly affected by the surrounding whey matrix. The XRD pattern of MMT exhibited one characteristic peak at 5.52°, representing the basal spacing of 1.60 nm, as is expected for calcium montmorillonite [51]. For the WPF/MMT nanocomposite, the peak has shifted to 4.65° (*i.e.* a basal spacing of 1.90 nm) and has broadened, indicating that the whey protein matrix is intercalated into the MMT interlayer space, leading potentially to a higher degree of disorder.

**Figure 4:**
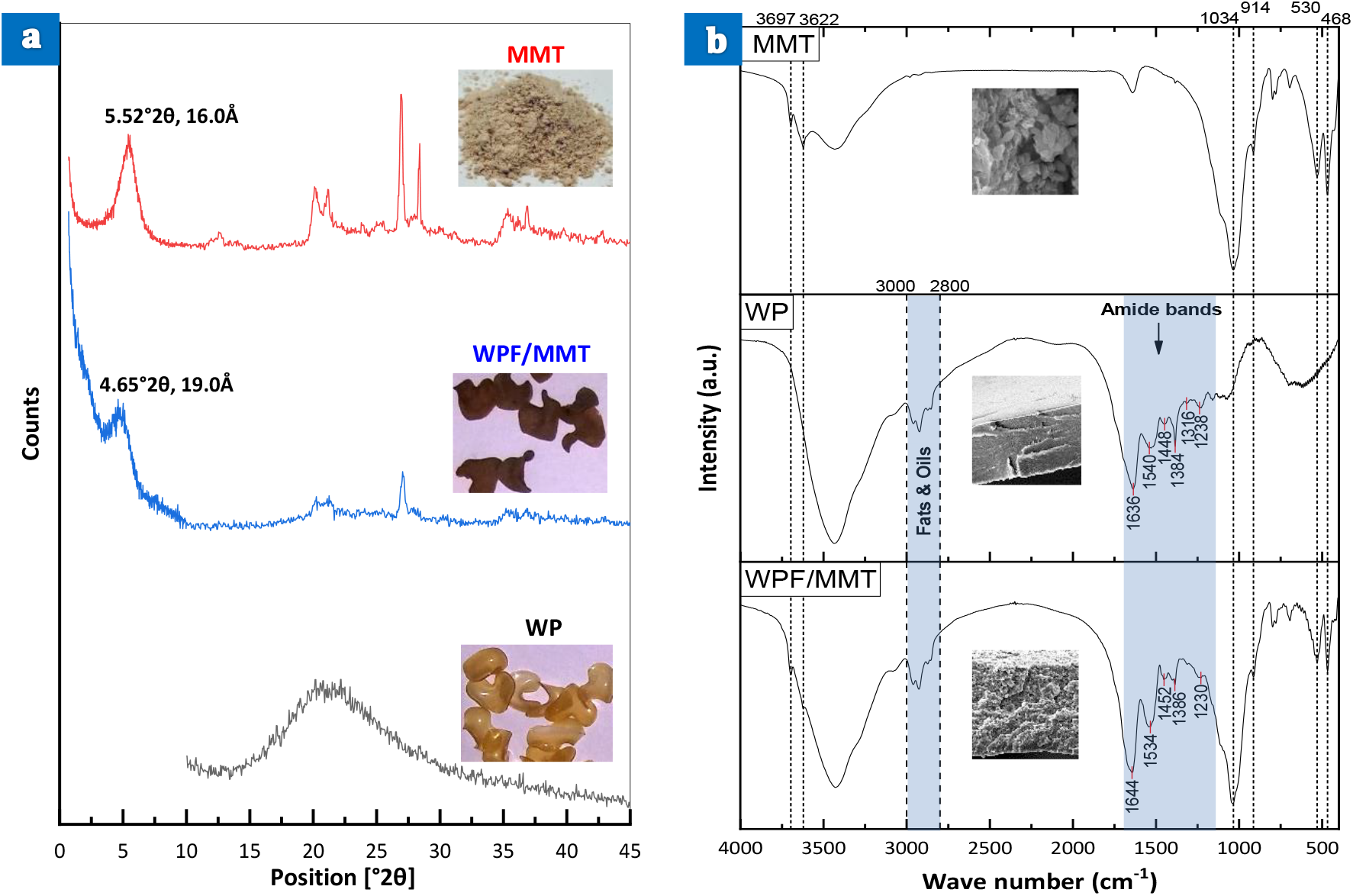
**(a)** XRD pattern of pristine MMT, WP, and WPF/MMT. *Insets:* macrostructures. **(b)** FT-IR spectrum of pristine MMT, WP, and WPF/MMT. Shaded areas indicate the characteristic peaks of fats and oils as well as proteins. Main peaks of MMT are found within the three pairs of stippled lines (wavenumbers are provided at the top of the graph). *Insets:* SEM structures.

#### 4.2.3. FT-IR Spectroscopy

The functional groups of the prepared nanocomposite were investigated by FT-IR analysis (**Fig. 4b**). The main IR absorption of the protein backbone is related to amide bands in the 1700-1200 cm^−1^ region [52] which can all be detected in both WP and WPF/MMT spectra (right shaded area in **Fig. 4b**). Also, there are some peaks around 2800-3000 cm^−1^ which can be assigned to fats and oils [53] in both WP and WPF/MMT spectra (left shaded area). Thus peaks between 2854 and 2923 are specific for dairy fatty acids. Furthermore, characteristic peaks of MMT (shown by grids) were observed in the composite too. As is listed in **Supp. Table 2**, WPF/MMT exhibited both MMT and WP main functional groups. However, in the composite, bands at 3697 and 3622 cm^−1^, related to free OH vibration, were reduced in intensity, implying that there are fewer free OH bands in the nanocomposite. Some of the free hydroxyl groups found in MMT may have formed hydrogen bonds with the whey proteins [54]. Furthermore, based on the FT-IR spectra of the WP/MMT and WPF/MMT presented in **Supp. Fig. 3** in the absence of nanofibrils, some peaks related to MMT and amide bands (shown by red and black stippled lines, respectively) are less intense, while changes in other peaks (indicated by arrows) show alterations in aluminosilicate functional groups. In contrast, amide band peaks around 1642, 1536, and 1448 cm^−1^ increased in intensity along with the peak around 3424 cm^−1^, indicating more hydrogen bonding in this composite. Note that after soaking in water, the WP films do not show any of the typical peaks of glycerol (five peaks in region 800 cm^−1^ to 1150 cm^−1^) [21]. This means that glycerol has been removed during soaking in water (*i.e.* purification step in **Fig. 1**).

#### 4.2.4. BET Analysis

BET analysis is a common technique for determining the surface area, possible adsorption mechanism, and pore shape of the porous materials, through measurement of the amount of N_2_ adsorption/desorption at 77 K [55]. The N_2_ sorption isotherm of WPF/MMT nanocomposite (**Fig. 5a**) curves upwards with increasing pressure and does not show a point B (*i.e.* a knee on the isotherm) which would otherwise indicate complete monolayer coverage. Therefore there is no recognizable monolayer formation. Moreover, the value of the C parameter (1.74) is below 2. These observations all indicate that the isotherm is type III, in which the interaction between adsorption and adsorbate is relatively weak, and the adsorbed molecules are clustered around favorable sites [55, 56]. The sorption isotherm showed a hysteresis loop which according to IUPAC nomenclature can be classified as Type H3 (no adsorption limit at high p/p_0_) which is in line with other clay composites [51, 57]. Type H3 loop is commonly seen in non-rigid combinations of plate-like particles (*e.g.*, clays) [55, 58].

**Figure 5:**
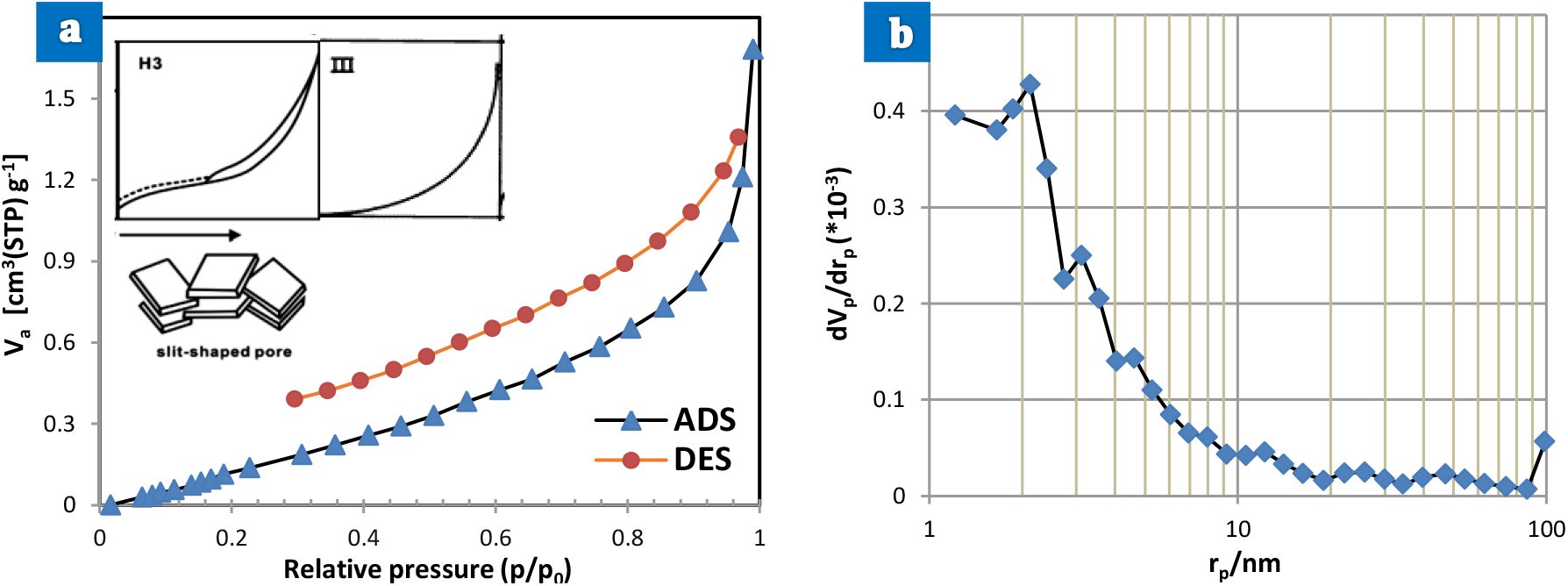
**(a)** Nitrogen adsorption/desorption isotherms (BET analysis) of the WPF/MMT Nanocomposite. Inset pictures show type III isotherm model (right) and hysteresis loop type H3 and the related pore shape (*i.e.* slit-shaped) (left) [59], and **(b)** corresponding BJH pore size distributions.

The specific surface area and pore radius were determined by the BJH method (**Fig. 5b**) to be 1.28 m^2^/g and 2.13 nm, respectively. It should be noted that the H3 hysteresis loop does not generally provide a reliable estimate of the pore size distribution [51, 56]. Besides, the nanocomposite sample used for the BET test was in flakes, not in powder form. The surface area of the composite is very modest in comparison with pristine MMT (220-270 m^2^/g), bentonite micropowder (67 m^2^/g) [60], or similar composites *e.g.* clay-cellulose composite (31.16 m^2^/g) [57], *Moringa oleifera* seed protein-montmorillonite composite (4.90 m^2^/g) [61] or carbon/montmorillonite nanocomposite (39.5 m^2^/g) [51].

### 4.3. Batch Dye Adsorption Studies

#### 4.3.1. Dye Removal Rate of Different Classes of Azo Dyes

The dye removal capability of WPF/MMT nanocomposite was examined using the eight azo dyes mentioned in **Supp. Table 1**. Cationic dyes (Chrysoidine-G and Bismarck brown-R) adsorbed much faster than the other dyes (**Fig. 6a**).

**Figure 6:**
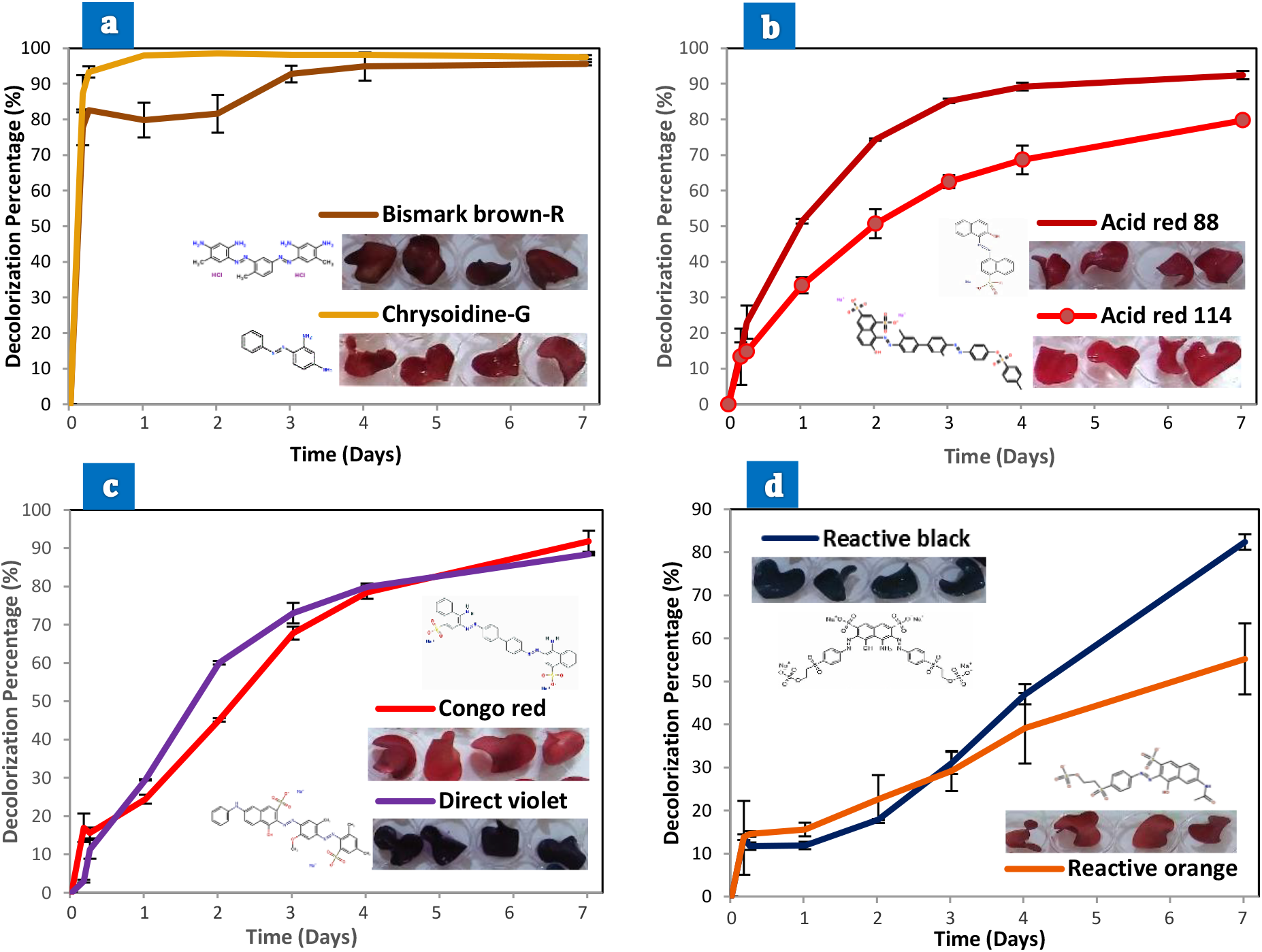
Adsorption of different azo dyes over time onto a 2% w/v WPF/MMT nanocomposite **(a)** Two cationic dyes, **(b)** two acid dyes, **(c)** two direct dyes, and **(d)** two reactive dyes. In all cases, the initial dye concentration was 250 mg/L. Inset photographs show the appearance of WPF/MMT after 1 week soaked in the dye solutions. The chemical structure of the dyes is also shown.

Most likely the negative charge of MMT helps adsorb the cationic dyes via ion exchange and electrostatic interactions. As regards acidic dyes, acid red 88 adsorbed faster (89% after 4 days) than acid red 114 (80% after 7days), which we ascribe to its smaller molecular structure (**Fig. 6b**). Congo red and direct violet 51 presented highly similar patterns in their adsorption timeline; both are ~80% adsorbed after 4 days but have not reached a plateau after a week (**Fig. 6c**). Finally, reactive dyes (reactive black 5 and reactive orange 16) adsorbed more slowly than the other dyes and required more time to reach equilibrium (**Fig. 6d**).

The dye removal timeline using WP and WP/MMT nanocomposite is presented in **Supp. Fig. 4.** In the first days, these two nanocomposites adsorbed Congo red, direct violet, and acid red 88 dyes significantly faster than WPF/MMT. However, over a longer time scale, the extent of adsorption on WP and WP/MMT decreased slightly for some dyes (*i.e.* acid red 88, direct violet 51, reactive orange and reactive black). We observed that WP and WP/MMT were not as stable as WPF/MMT over a week of incubation with the dyes. Also, they become more swollen than the amyloid-containing composite, WPF/MMT (**Supp. Fig. 5**). Swelling depends on the solvent and the cross-link density [62, 63]. Thus, in the presence of nanofibrils the composite is more cross-linked, and hence reinforces the nanocomposite [23], making it more stable in different dye solutions.

We note that to achieve this stability, drying the nanocomposite at 100°C, for 1 h (the last stage in **Fig. 1**) is necessary as heat treatment, to form enough cross-link between nanofibrils and MMT components (see sketch in **Fig. 3** and **Supp. Fig. 5d**).

#### 4.3.2. Effect of Initial Dye Concentration

We decided to carry out a more detailed analysis of the absorption process using the cationic dye Chrysoidine-G, which showed the fastest and most extensive level of absorption. **Fig. 7a** shows the influence of the different initial dye concentrations (from 25 to 250 mg/L) and contact time on Chrysoidine-G dye removal, using WPF/MMT (8 g/L).

**Figure 7:**
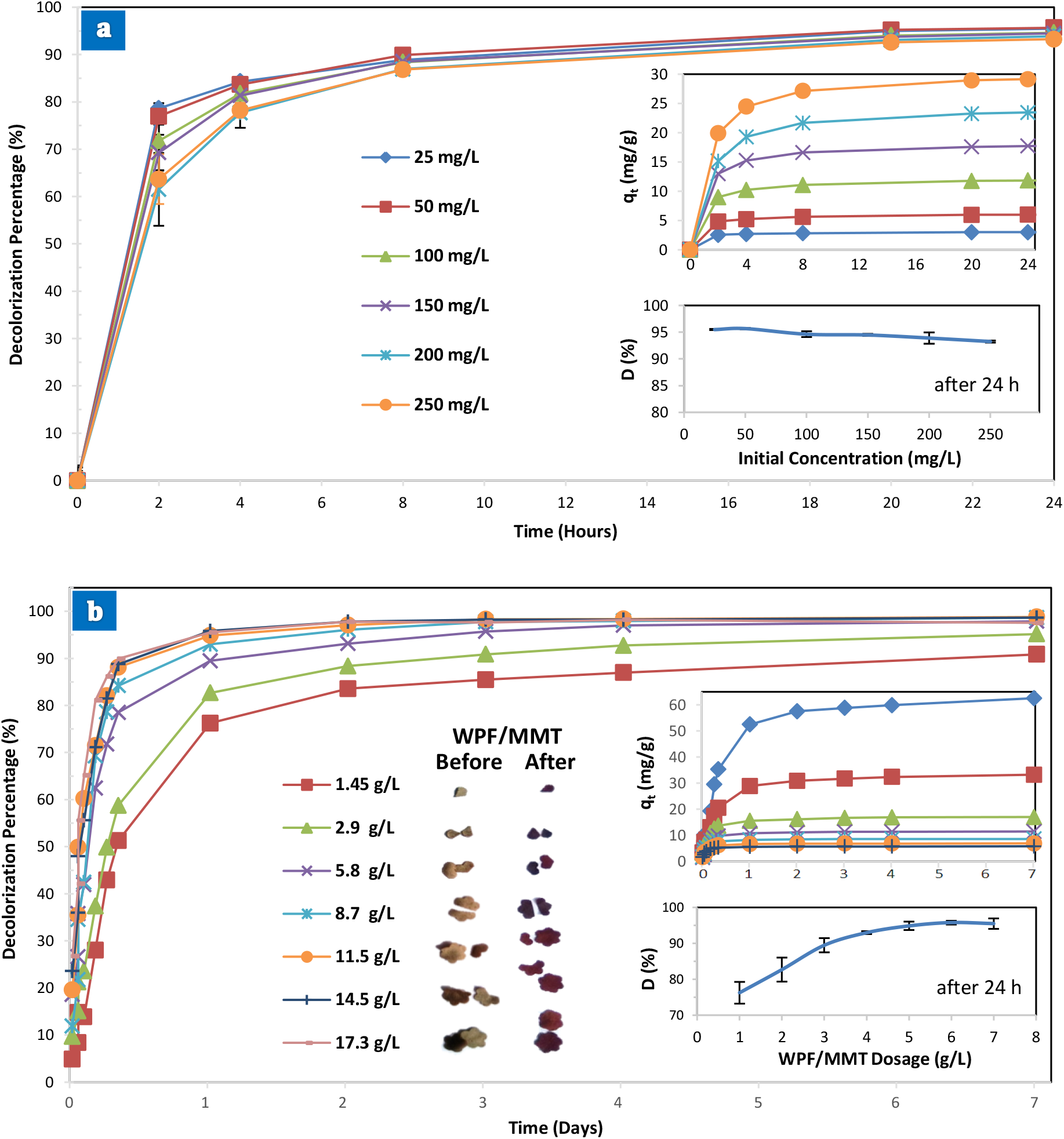
Effect of **(a)** dye concentration (using 8 g/L adsorbent), and **(b)** WPF/MMT dosage (using 100 mg/L dye) on Chrysoidine-G dye decolorization percentage, at 35 °C. Amount of dye adsorbed by WPF/MMT (q_t_) (upper inset graph) and the effect of WPF/MMT dosage (g/L) on adsorption percentage (lower inset graph). Inset pictures show WPF/MMT before and after the adsorption process.

Although the rate of adsorption (*i.e.* the plot slope) decreased slightly with increasing initial concentration (**Fig. 7a**), the equilibrium adsorption percentage did not change very much, and WPF/MMT was able to adsorb more than 93% of the dye at all concentrations over a 24-h period. In contrast, for most adsorbents, a decrease in adsorption percentage has been reported with increasing initial dye concentration [64–66]; moreover, the amount of the adsorbed dye (q_t_,) is proportional to the increase in initial dye concentration, as when the initial dye concentration was doubled, q_t_ was doubled too (R^2^≥0.9997) (**Fig. 7a**, upper inset graph). Thus, the similar relative absorption at different dye concentrations indicates that the only limiting factor, in this adsorption dose, is the bulk diffusion of dye molecules from the solution to the boundary layer on the adsorbent surface [46]. These results are consistent with the results of BET, isotherms, and kinetics studies (see below), which indicate the existence of an unlimited number of adsorption sites under the present conditions (*i.e.* we do not reach the full capacity of the adsorbent under our experimental conditions).

#### 4.3.3. Effect of Adsorbent Dose

The optimum WPF/MMT dose for adsorbing Chrysoidine-G dye (100 mg/L) was investigated using different WPF/MMT doses over a week (**Fig. 7b**). Increasing the WPF/MMT dose from 1.45 to 5.8 g/L increased decolorization rate; however, further dose increase does not significantly affect either decolorization or q_e_. Therefore, approximately 9 g/L is the optimum adsorbent dosage for the adsorption of Chrysoidine-G by WPF/MMT.

#### 4.3.4. Effect of Initial Dye pH

The solution pH influence on Chrysoidine-G (100 mg/L) adsorption using WPF/MMT was studied at a wide pH range of 2-10. The adsorption rate at pH≥ 6 is almost constant at 90%, except the pH= 9 which shows the minimum adsorption rate (**Fig. 8a**). However, the maximum adsorption (> 98%) occurred in acidic pH, less than or equal to 5, where a color change is also visible (see inset image in **Fig. 8a**). This suggests that at pH≤ 5, the prevalent form of Chrysoidine-G is protonated (*i.e.* amino groups are positively charged), and can be adsorbed more efficiently by negatively charged WPF/MMT. While, at pH≥ 6 the neutral form of Chrysoidine-G is dominant; accordingly, adsorption can proceed via other interactions (*e.g.* hydrophobic and hydrogen forces) rather than electrostatic adsorption.

**Figure 8:**
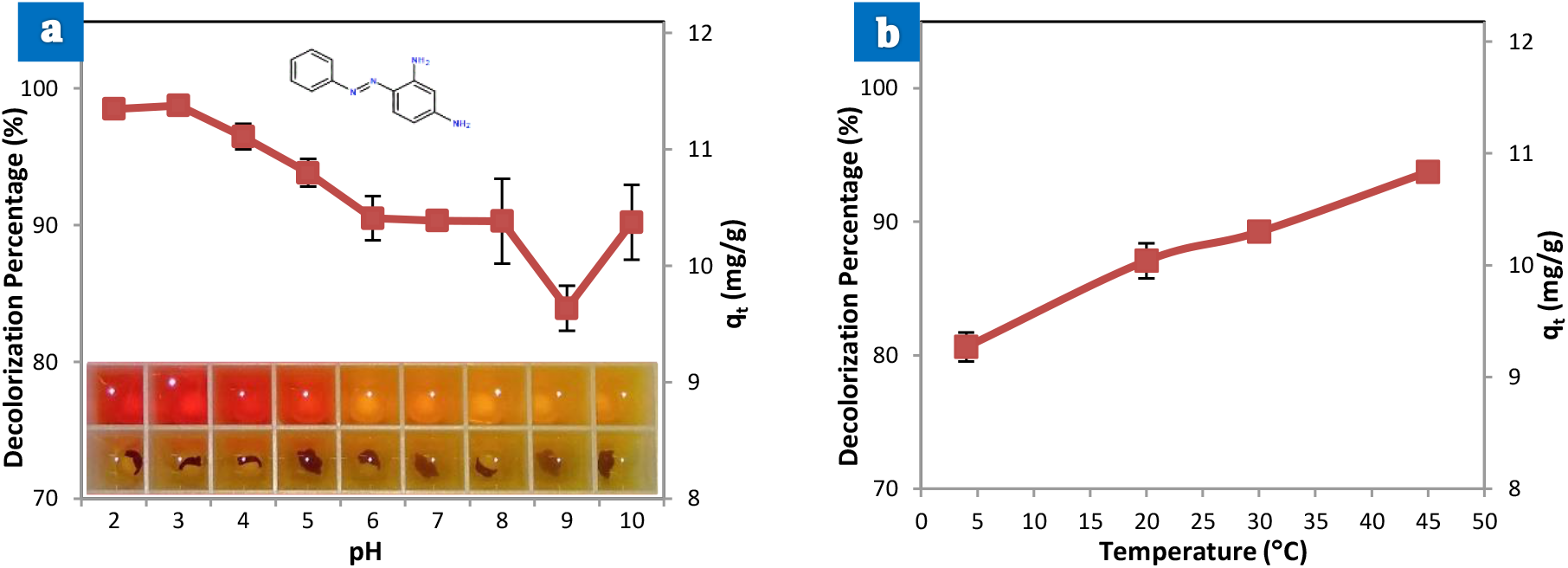
Effect of **(a)** pH, and **(b)** temperature on the adsorption of Chrysoidine-G dye (100 mg/L) on WPF/MMT (8.7 g/L). pH study analyzed at 37 °C, after 2 days, and Temperature study after 3 days. *Inset*: dye solutions with different pH; top-row: control and bottom-row: with WPF/MMT, after 2 days.

#### 4.3.5. Effect of Temperature

The influence of temperature on Chrysoidine-G (100 mg/L) adsorption was studied using WPF/MMT at 4, 20, 30, and 45°C (**Fig. 8b**). It was observed that the adsorption rate and q_e_ increased with an increase in the temperature. The diffusion of the dye molecules increases with a rise in the temperature leading to a higher mass transform rate from the bulk solution to the boundary layer on the WPF/MMT [46]. This result is in agreement with the study of Purkait et al. (2010) on Chrysoidine adsorption using activated carbon [67] and is compatible with the kinetics results suggesting chemisorption and surface sorption as the main rate-limiting factors (see section 4.5).

### 4.4. Adsorption Isotherms

Adsorption isotherms studies provide useful information on how solutes interact with adsorbents, and it is essential in optimizing the use of adsorbents. In this study, equilibrium adsorption parameters of Chrysoidine-G adsorption onto WPF/MMT were investigated at 37°C using different adsorbent dosages and subsequently fitted with linear (Nernst), Longmuir, Freundlich, Temkin, and D-R Isotherm models.

The equilibrium adsorption data showed a simple linear relationship between the total and adsorbed amount of dye (**Fig. 9a**). Similarly, the best-fitted curves (R^2^> 957) obtained using the equations describing the Langmuir, Freundlich, and Temkin isotherms all approach a linear form (**Fig. 9a** and summarized in **Table 1**). Accordingly, the simplest possible equation (with the least number of parameters) should be selected, *i.e.* the simple linear (*i.e.* Nernst or Henry) isotherm. In a model with a linear isotherm (C-class), the number of adsorption sites remains constant until saturation of the adsorbent. This means that the adsorption surface increases during the adsorption process, and can be considered as an unlimited number of adsorption sites because as soon as each molecule is adsorbed, a new vacant site is generated. **Fig. 9b** illustrates the conditions of this C-class system; the adsorbed molecules penetrate the inner layers of the adsorbent, and since the entrance of pores is not occupied, the adsorption process proceeds continuously [68]. A microscopic look at the boundary layer of WPF/MMT (**Fig. 9c**), reveals that the adsorbent surface in the solution exhibited a gel-like surface where dye molecules can penetrate, and aggregate in some, probably more favorable, places, as indicated with black arrows in the picture. Thus, the process proceeds continuously. This conforms to the BET results in section 4.2.4 which follows the type III isotherm model, indicating that the adsorbed molecules are clustered around favorable sites, and the adsorption process can proceed continuously.

**Figure 9:**
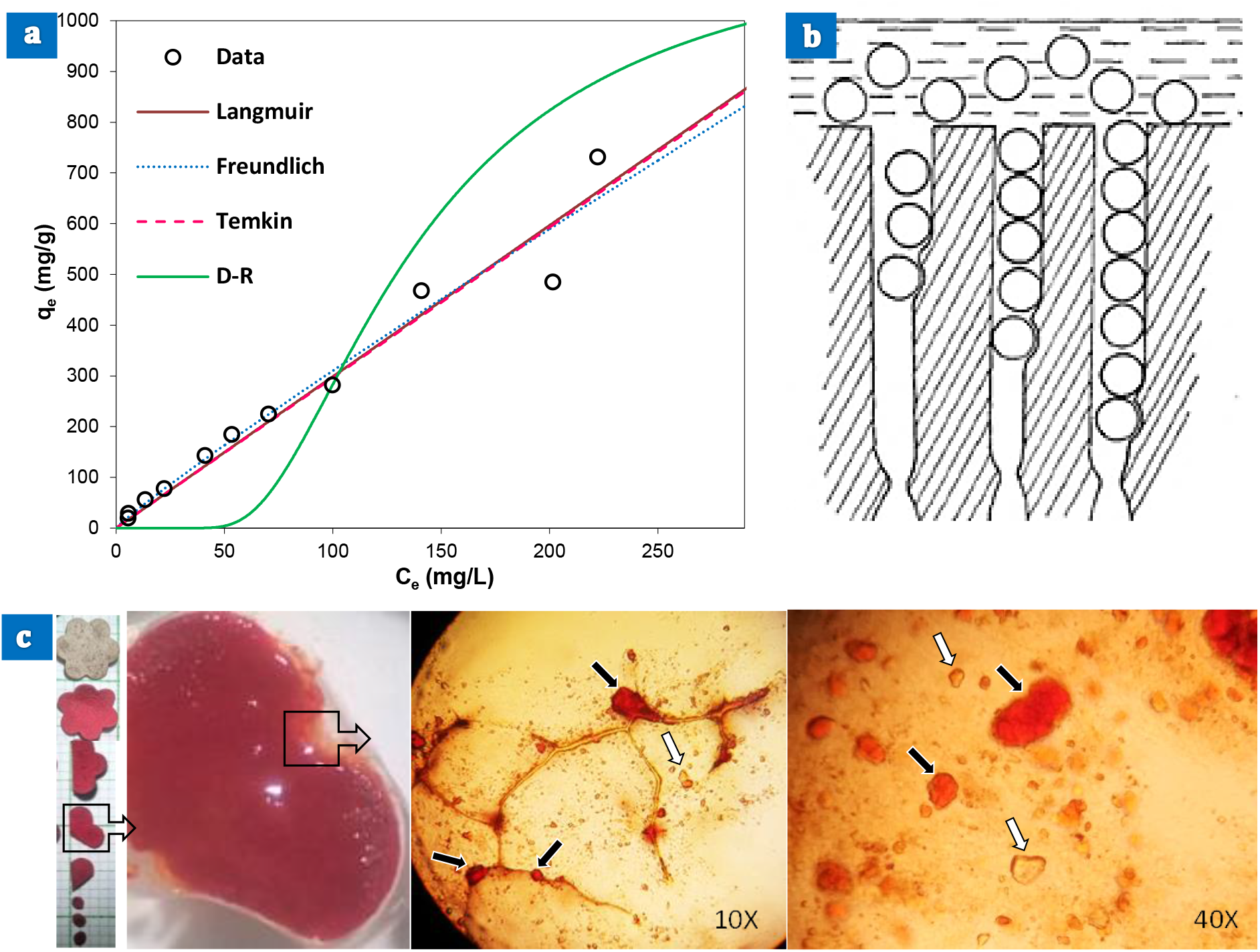
(**a**) Nonlinear adsorption isotherms of Chrysoidine-G onto different doses of WPF/MMT at 37°C. **(b)** Scheme illustrating linear isotherm (Giles, Smith, & Huitson, 1974; Giles, Smith, Huitson, & science, 1974), **(c)** microstructure of WPF/MMT after adsorption of Chrysoidine-G, Favorable places for dye aggregation: *black arrows*, and unfavorable places: *white arrows*

**Table 1:** Comparison between linear and non-linear forms of adsorption isotherms for Chrysoidine-G onto WPF/MMT at 37°C

|  | Linear (Nernst) | Langmuir | Freundlich | Temkin | D-R |
| --- | --- | --- | --- | --- | --- |
| Non-linear form | - | 0.9571 | 0.9593 | 0.9570 | 0.5441 |
| Linear form | 0.9594 | 0.9501 | 0.9891 | 0.7842 | 0.6872 |

To gain a better insight into the adsorption mechanism, we also studied the linearized versions of the adsorption isotherms (**Supp. Fig. 6**). Comparing the related R^2^ amounts of these two forms (**Table 1**), it is noticeable that Freundlich fits the adsorption data better than Langmuir model. This agrees with the BET results (**Fig. 5a**) which show no completion of monolayer adsorption.

### 4.5. Kinetics of Adsorption

To study the mechanism of the process, kinetics of Chrysoidine-G adsorption onto WPF/MMT were analyzed using pseudo-first-order, pseudo-second-order, and intra-particle diffusion models, based on the raw adsorption data presented in **Fig. 10a**.

**Figure 10:**
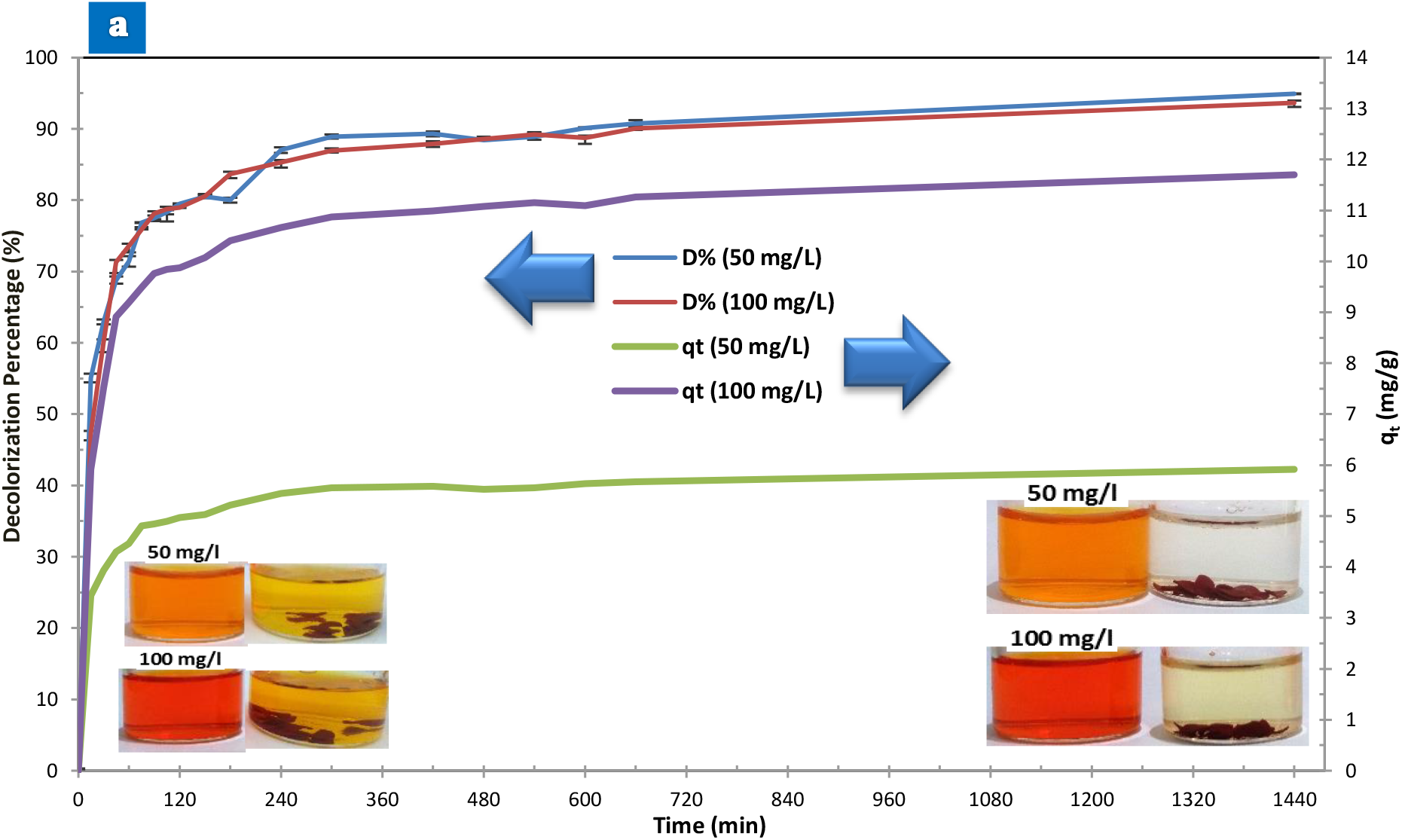
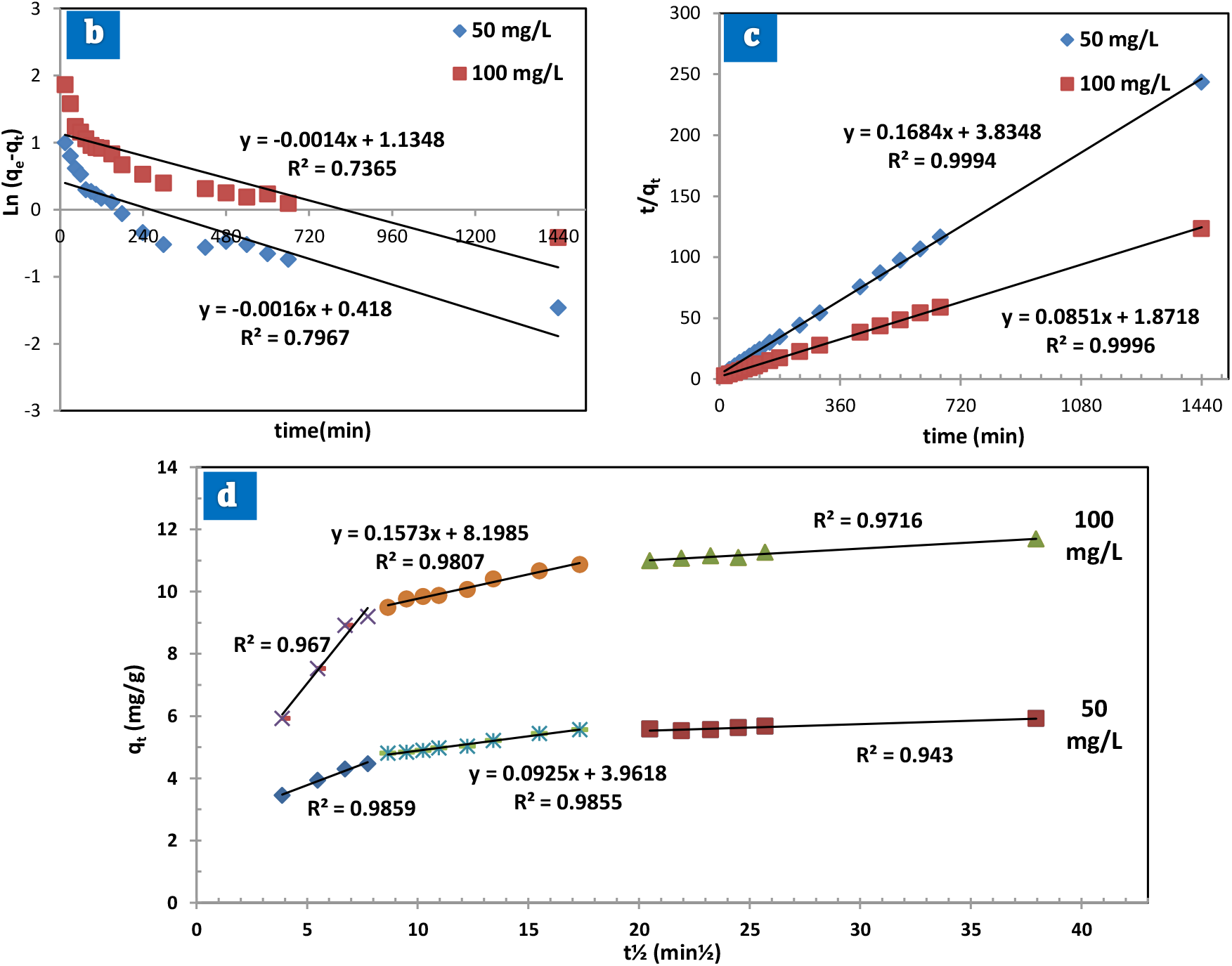
Kinetic study of Chrysoidine-G dye adsorption onto WPF/MMT (8 mg/L) at 35°C over 24 h, in a volume of 10 mL. **(a)** Decolorization percentage and amount of dye adsorbed by WPF/MMT (q_e_), inset pictures show Chrysoidine-G 50 and 100 mg/L without and with WPF/MMT after 80 minutes (left), and after 48 hours (right). **(b)** Pseudo-first-order model, **(c)** pseudo-second-order model, **(d)** intra-particle diffusion model for Chrysoidine-G adsorption onto WPF/MMT.

#### Pseudo-first-order

The pseudo-first-order parameters for the adsorption using WPF/MMT are listed in **Table 2**. It is clear from **Fig. 10b**, that the pseudo-first-order equation provides an extremely poor fit to the data. Furthermore, the calculated q_e_ is not in agreement with the experimental q_e_ (even after iterating altering the q_e_). We conclude that experimental data is not fitted well with this model.

**Table 2:** Kinetic parameters for Chrysoidine-G adsorption onto WPF/MMT (8 mg/L) at 35°C

| $C_0$<br>(mg/L) | $q_e(\text{exp})$<br>(mg/g) | Pseudo-first-order | | | Pseudo-second-order | | | Intra-particle diffusion | | |
| --- | --- | --- | --- | --- | --- | --- | --- | --- | --- | --- |
| | | $K_1$<br>(1/min) | $q_e(\text{cal})$<br>(mg/g) | $R^2$ | $K_2$<br>(g/mg min) | $q_e(\text{cal})$<br>(mg/g) | $R^2$ | $K_i$<br>(mg/g min<sup>1/2</sup>) | $C_i$<br>(mg/g) | $R^2$ |
| 50 | 5.67 | -0.0016 | 1.52 | 0.7967 | 0.0074 | 5.94 | 0.9994 | 0.0925 | 3.9618 | 0.9855 |
| 100 | 11.26 | -0.0014 | 3.11 | 0.7365 | 0.0039 | 11.75 | 0.9996 | 0.1573 | 8.1985 | 0.9807 |

#### Pseudo-second-order

The pseudo-second-order parameters are presented in **Table 2**. Clearly, this leads to a much better fit than the pseudo-first-order results, as well as a good agreement between experimental (*i.e.* measured directly from the experiment) and calculated q_e_ (*i.e.* calculated from the plot slope (**Fig. 10c**), eq. 10), making it clear that this is an excellent model for Chrysoidine-G adsorption. As a corollary, this implies that chemisorption is the rate-limiting step of Chrysoidine-G adsorption using WPF/MMT [45]. Also, it can be observed that the value of the rate constant k_2_ is halved when the initial dye concentration is doubled. Thus Chrysoidine-G adsorption on WPF/MMT depends almost only on the diffusion and is not limited by the amount of adsorbent, indicating the presence of increasing or infinitive adsorption sites. This is in agreement with the results of our adsorption isotherms study in section 4.4 which has been mentioned that the adsorption of each molecule generates a new free site (**Fig. 9b**).

#### Intra-particle diffusion

The intra-particle diffusion parameters are shown in **Table 2**. As shown in **Fig. 10d**, the intra-particle diffusion curve (adsorption versus square root of time, eq. 11) shows three linear regions, revealing three stages in the adsorption process. A similar three-stage plot has been reported on adsorption of malachite green by cellulose nanofibril aerogel [66], or chitin hydrogels [69], adsorption of multi-azo dye using chitosan-starch hydrogel [70], and Congo red adsorption by cashew nutshell [64].

According to the interpretation of this three-stage process [66, 69–71], the first stage (up to 60 minutes, 70% adsorption) is related to the diffusion of dye molecules from the bulk solution to the external surface of WPF/MMT. The second stage (up to 5 h, ~90% adsorption) can be attributed to intra-particle diffusion of dye molecules into the interior of WPF/MMT, and finally, the third linear stage is related to the equilibrium phase of the adsorption process and reflects the decrease in dye concentration [69, 72]. Thus, it can be concluded that the adsorption data can be described with the intra-particle diffusion model. The constant C_i_, which is related to the boundary layer thickness, increases when the dye concentration increases. Indicating an increase in the thickness of the boundary layer with increasing the initial concentration, therefore the rule of surface sorption becomes more important as the rate-limiting step [64].

Most of our experiments were performed in a small volume (1.5 mL). To test for possible scale-ups, the kinetic experiment was performed on a larger scale of 10 mL (**Fig. 10a**). Comparison of the results of adsorption of Chrysoidine-G (100 mg/L) using 8 g/L of WPF/MMT shows that the dye removal percentage and q_t_ values are very close in both volumes; this suggests that the adsorption process may be scaled up without significant changes in the results.

## 5. Conclusions

In this study, an easy handling bio-nanocomposite was developed from WPC using proteinaceous nanofibrils together with MMT (WPF/MMT) with simple and convenient separation. The structure of the resulted nanocomposite was characterized using SEM, FT-IR, BET, and XRD which indicated WPF/MMT is a nanocomposite with intercalated and exfoliated structure. Investigating the adsorption efficacy for the different classes of azo dyes with different physicochemical properties revealed that WP, WP/MMT, and WPF/MMT can be used to adsorb a wide range of azo dyes. WPF/MMT presented a better performance in adsorbing cationic dye. Although WP and WP/MMT adsorbed acid, direct and reactive dyes more rapidly than WPF/MMT, only WPF/MMT remained stable in all studies for a long time, probably because of the presence of nanofibrils, higher cross-link density, and lower swelling capacity. In further studies on the decolorization process, it was determined that the adsorption phenomenon was almost independent of the dye concentration, and the optimum adsorbent dosage was ~9 g/L. To decipher the mechanism underlying the adsorption process, different isotherm and kinetic models were investigated. Since kinetic data could be fitted very well using a pseudo-second-order model, we conclude that chemisorption plays a rate-limiting role in the adsorption process. Moreover, comparing linear and non-linear forms of adsorption isotherms suggests that the process is mostly fitted with the linear (Nernst) isotherm model, indicating the existence of unlimited absorption sites, which in turn conforms with the results of BET (Type III isotherm), kinetic studies (both pseudo-second-order and intra-particle diffusion), and also optical microscopy. Overall, whey protein-based polymer and nanocomposites have clear potential as effective and easy-separable adsorbents for different classes of dyes or even other pollutants from wastewater.

## Acknowledgments

This work was supported by the Bioprocess Engineering Department, Institute of Industrial and Environmental Biotechnology, National Institute of Genetic Engineering and Biotechnology; and Center for International Scientific Studies & Collaboration (CISSC), Ministry of Science Research and Technology. D.E.O. is grateful for support from the Independent Research Foundation Denmark | Natural Sciences (grant no. 8021-00208B). The authors would like to thank Farhang Aliakbari for helpful discussions, and Zahra Najarzadeh for help with TEM images. Tayebe Bagheri Lotfabad is acknowledged for helpful suggestions in the film preparation process.

