## Supplementary material for "Adsorption of Azo Dyes by a Novel Bio-Nanocomposite Based on Whey Protein Nanofibrils and Nano-clay: Equilibrium Isotherm and Kinetic Modeling": Supplementary.pdf

**Table S1:** Molecular weight, maximum absorbance wavelength (nm), and structure of the dyes used in this study.

| CHRYSOIDINE-G | | BISMARCK BROWN-R | |
|---|---|---|---|
| MW= 248.7 Da, MonoAzo | 449 nm | MW= 461.4 Da, DiAzo | 468 nm |
| 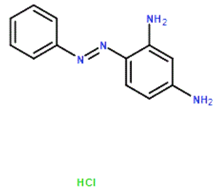 | | 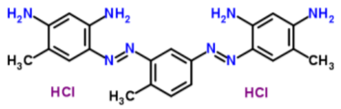 | |
| **CONGO RED** | | **DIRECT VIOLET 51** | |
| MW= 696.7 Da, MonoAzo | 500 nm | MW= 719.7 Da, DiAzo | 549 nm |
| 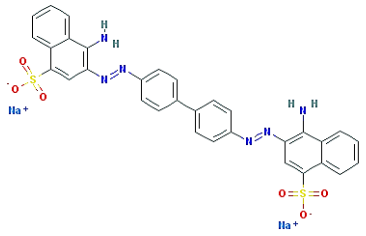 | | 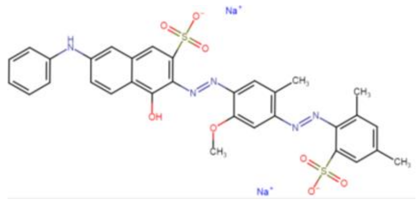 | |
| **ACID RED 88** | | **ACID RED 114** | |
| MW= 400.38 Da, MonoAzo | 505 nm | MW= 830.8 Da | 514 nm |
| 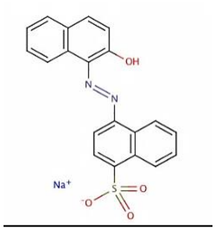 | | 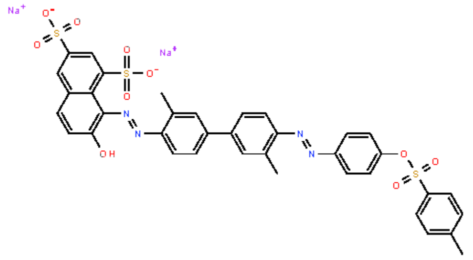 | |
| **REACTIVE ORANGE 16** | | **REACTIVE BLACK 5** | |
| MW= 617.5 Da, MonoAzo | 494 nm | MW=991.82 Da, DiAzo | 597 nm |
| 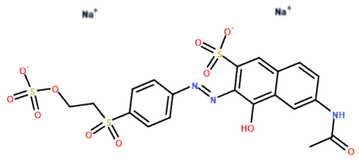 | | 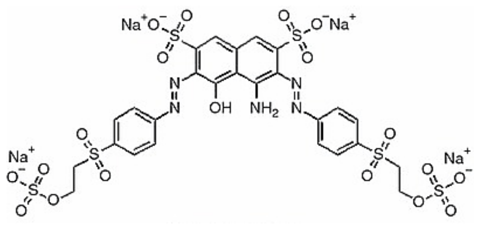 | |

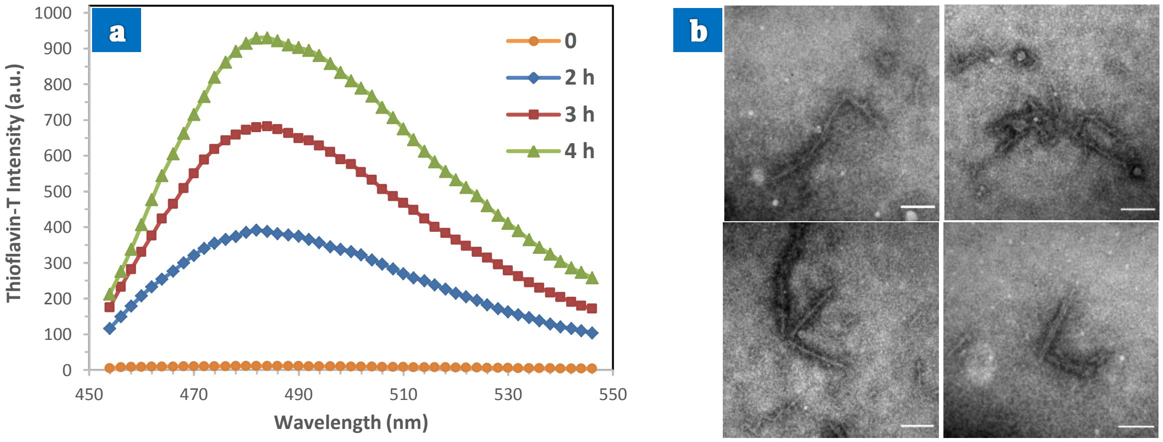


**Figure S1: Analyzing the formation of protein nanofibrils in the WP film-forming solution. (a)** Thioflavin-T fluorescence assay, samples were incubated at pH 2 and 85℃ **(b)** Overview of WP fibrils by TEM images, after fibrillation step and increase the pH to 7. Scale bars are 200 nm.

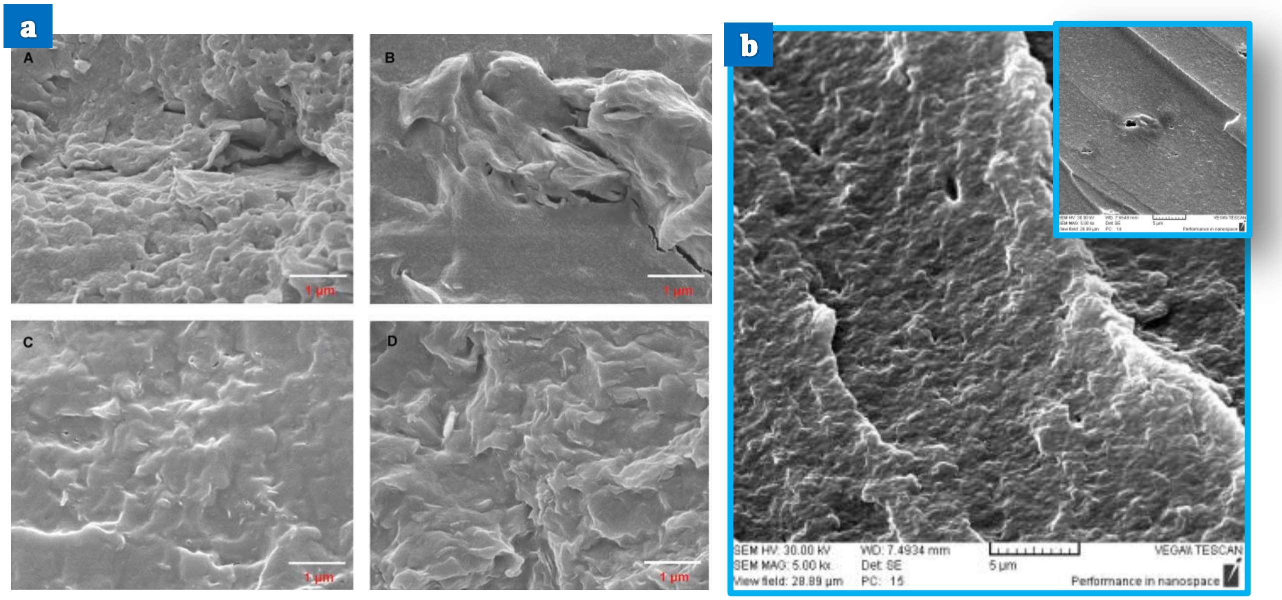


**Figure S2:** Comparison between SEM cross-sectional micrographs of **(a)** Kumar et al. soy protein isolate bio-nanocomposite films with 5%, 15% of modified MMT (Kumar, Sandeep, Alavi, Truong, & Gorga, 2010), adopted with permission. **(b)** Manually torn WP film in this study, the inset image shows WP film's fractured cross-section.

**Table S2:** FTIR characteristic peaks of WP, MMT, and WPF/MMT

| Functional Group | Vibration | Frequency Range ($cm^{-1}$) | Reference | WP | MMT | WPF/MMT |
|---|---|---|---|---|---|---|
| **Hydrogen Band** | O-H stretch | 3100-3600 | (Ben Faust, 2005) | ✓ | ✓ | ✓ |
| **Amide Bands** | Amide band I | 1700-1600 | (Kong & Yu, 2007) | 1636 | - | 1644 |
| | Amide band II | 1500-1400 | (Kong & Yu, 2007) | 1448 | - | 1452 |
| | Amide band III | 1400-1200 | (Kong & Yu, 2007) | 1384 | - | 1386 |
| **Fats and Oils (Dairy products)** | C-H stretch (carbonyl groups) | 2854-2923 | (Andrade et al., 2019) | ✓ | - | ✓ |
| **MMT** | Free O-H | 3697, 3622 | (Ben Faust, 2005) | - | ✓ | ✓ |
| | Si-O stretch | 1034, 695 | (Tong et al., 2018) | - | ✓ | ✓ |
| | Quartz | 798 | (Tong et al., 2018) | - | ✓ | ✓ |
| | Al-O-Si, Si-O-Si | 530, 468 | (Tong et al., 2018) | - | ✓ | ✓ |

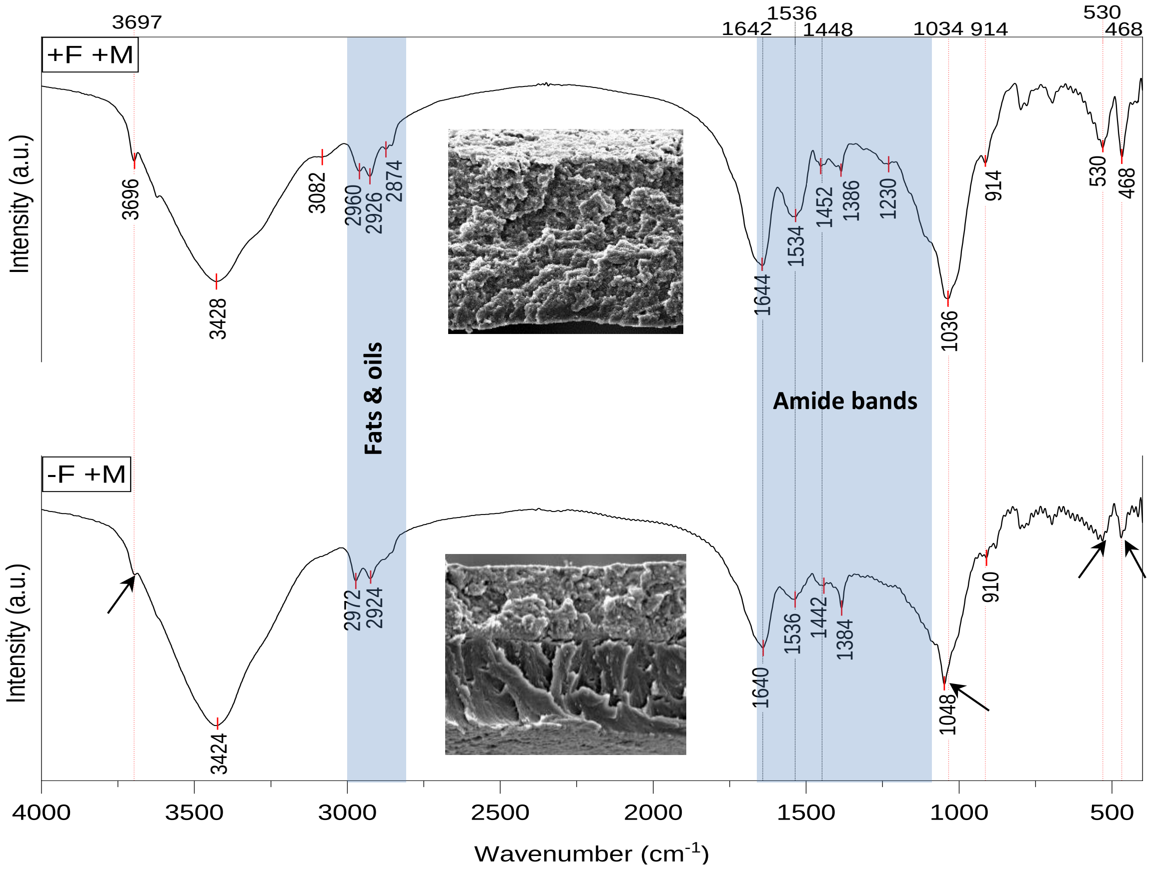


**Figure S3: FTIR spectrums of the WPF/MMT and WP/MMT nanocomposites.** stippled lines show the main peaks of MMT and shaded areas indicate the characteristic peaks of fats and oils as well as proteins. In WP/MMT (whitout amyloid fibrils), characteristic peaks correspond to Si-O-Si and Al-O-Si, as well as amide bands at around 1642, 1536 and 1448 $cm^{-1}$ are distinctively shorter than in WPF/MMT. In contrast, the peak around 3424 $cm^{-1}$ has been deeper indicating more hydrogen bands. Also, peak at 1036 $cm^{-1}$, corresponding to Si-O stretching vibration, shifted to 1048 $cm^{-1}$ and weakened, indicating a change in these functional groups in WP/MMT. Nevertheless, in WPF/MMT, peaks related to functional groups of MMT and WP present. *Insets:* SEM structures.

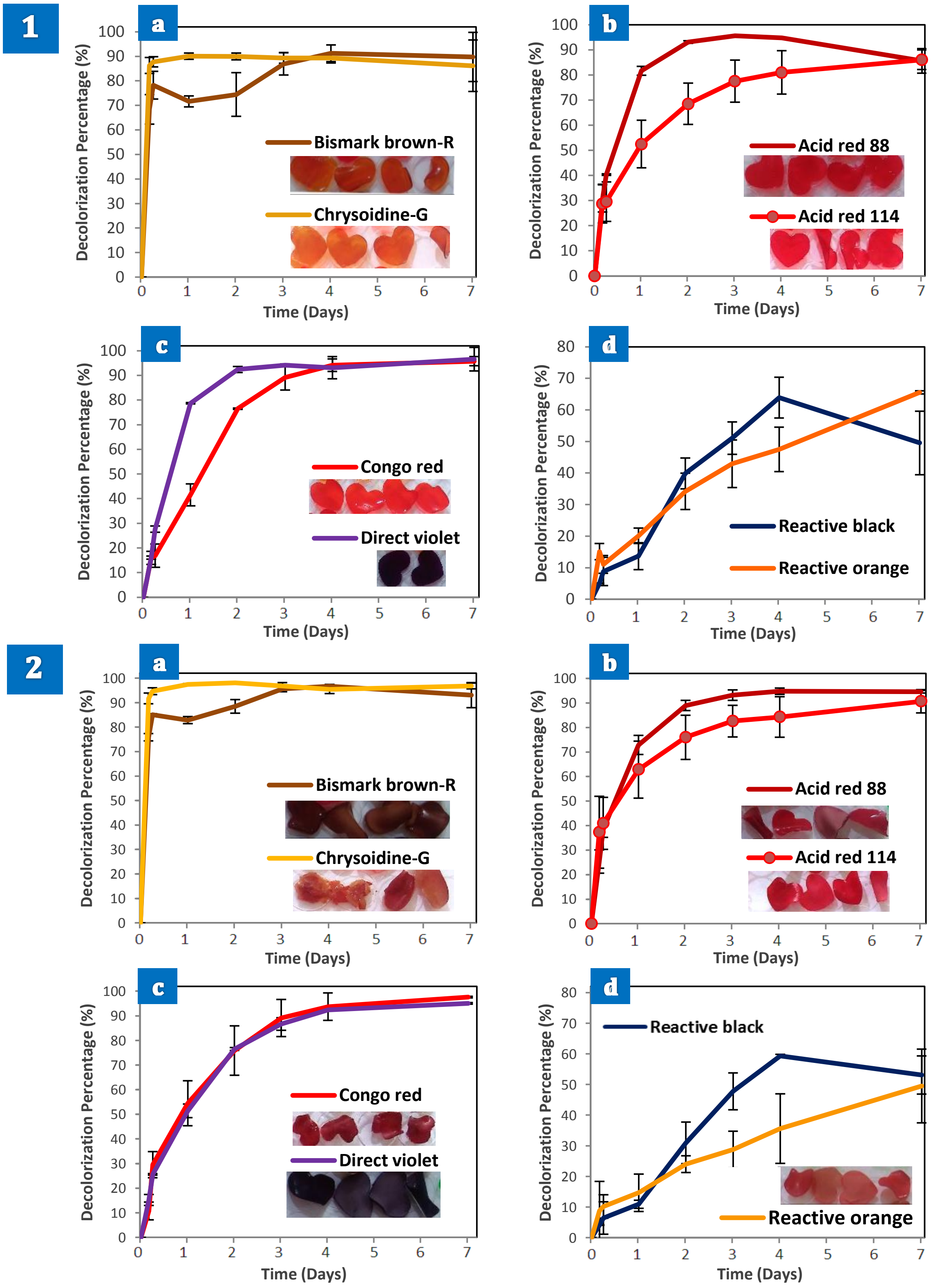


**Figure S4: Adsorption of different dyes over time onto (1) WP, and (2) WP/MMT**: **(a)** Two cationic dyes, **(b)** two acid dyes, **(c)** two direct dyes, and **(d)** two reactive dyes. In all cases, the initial dye concentration was 250 mg/L. Inset photographs show the appearance of WPF/MMT after 1 week soaked in the dye solutions. some of adsorbents are destroyed in some dyes' solution in a week and are absent in the graphs.

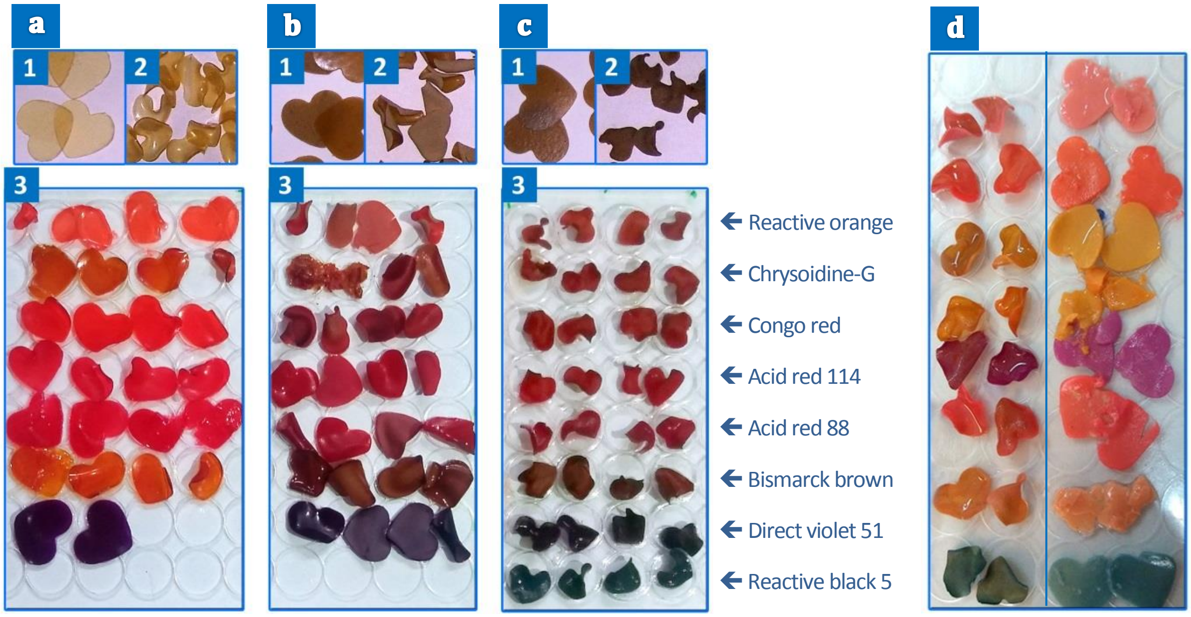


**Figure S5:** Photograph of **(a)** WP, **(b)** WP/MMT and **(c)** WPF/MMT: **(1)** before, and **(2)** after soaking in water and heat treatment, **(3)** show the adsorbents after immersing in the eight azo dye solutions for a week. WP and WP/MMT swelled more than the WPF/MMT, and some of them are destroyed in some dyes' solution in a week and are absent in the figure. **(d)** The effect of heat treatment (100°C, 1 h) on WPF/MMT after immersing in the dyes for 4 days. left: with heat treatment, right: without heat treatment.

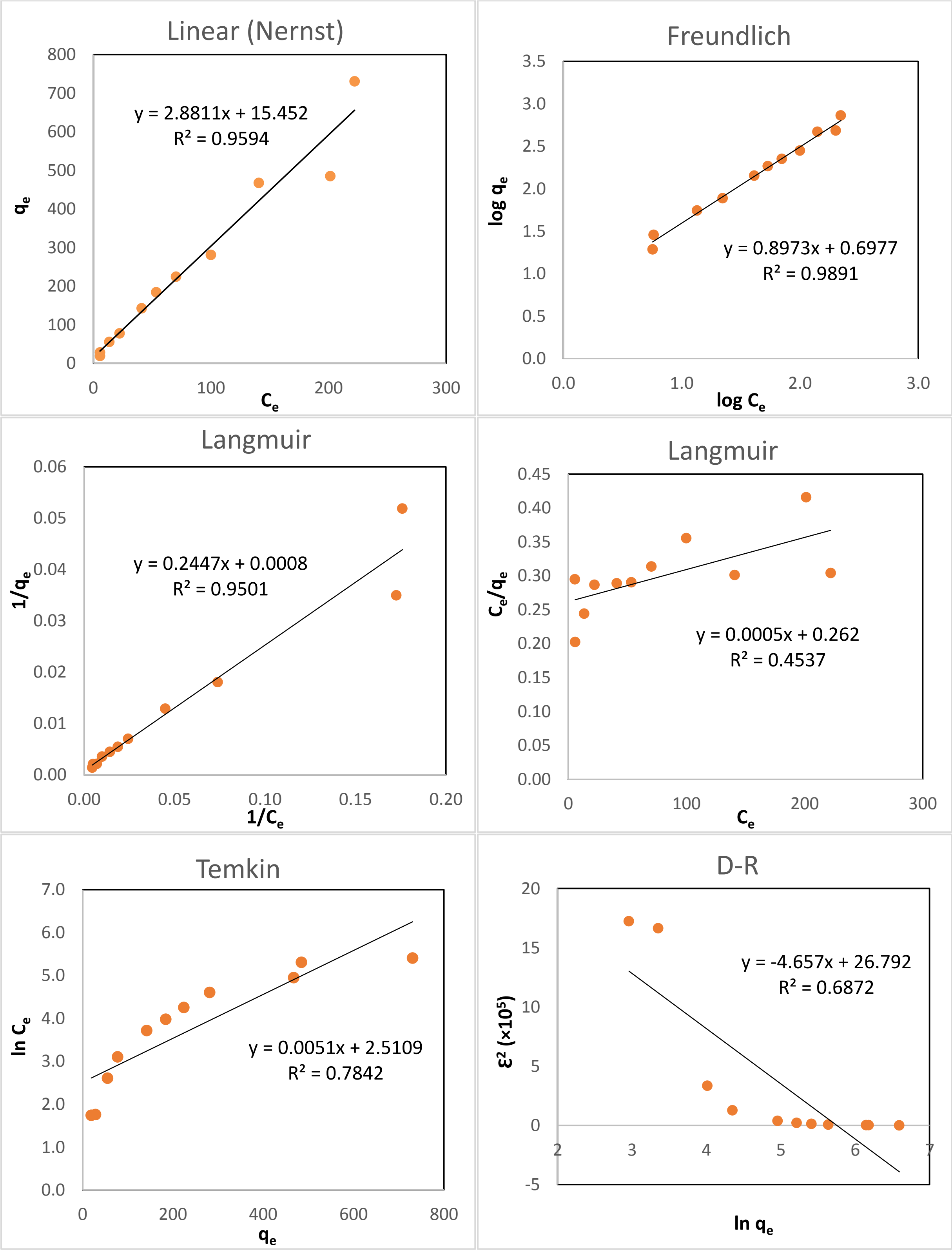


**Figure S6: Linear form of adsorption isotherms of Chrysoidine-G onto WPF/MMT at 37℃**
